# Tissue of Origin Dictates GOT1 Dependence and Confers Synthetic Lethality to Radiotherapy

**DOI:** 10.1101/714196

**Authors:** Barbara S. Nelson, Lin Lin, Daniel M. Kremer, Cristovão M. Sousa, Cecilia Cotta-Ramusino, Amy Myers, Johanna Ramos, Tina Gao, Ilya Kovalenko, Kari Wilder-Romans, Joseph Dresser, Mary Davis, Ho-Joon Lee, Zeribe C. Nwosu, Scott Campit, Oksana Mashadova, Brandon N. Nicolay, Zachary P. Tolstyka, Christopher J. Halbrook, Sriram Chandrasekaran, John M. Asara, Howard C. Crawford, Lewis C. Cantley, Alec C. Kimmelman, Daniel R. Wahl, Costas A. Lyssiotis

## Abstract

Metabolic programs in cancer cells are influenced by genotype and the tissue of origin. We have previously shown that central carbon metabolism is rewired in pancreatic ductal adenocarcinoma (PDA) to support proliferation through a glutamate oxaloacetate transaminase 1 (GOT1)-dependent pathway. Here we tested if tissue type impacted GOT1 dependence by comparing PDA and colorectal cancer (CRC) cell lines and tumor models of similar genotype. We found CRC to be insensitive to GOT1 inhibition, contrasting markedly with PDA, which exhibit profound growth inhibition upon GOT1 knockdown. Utilizing a combination of metabolomics strategies and computational modeling, we found that GOT1 inhibition disrupted glycolysis, nucleotide metabolism, and redox homeostasis in PDA but not CRC. These insights were leveraged in PDA, where we demonstrate that radiotherapy potently enhanced the effect of GOT1 inhibition on tumor growth. Taken together, these results illustrate the role of tissue type in dictating metabolic dependencies and provide new insights for targeting metabolism to treat PDA.

## Introduction

Metabolic processes are rewired in cancer to facilitate tumor survival and growth^1^. Accordingly, there is interest in defining the metabolic pathways utilized by cancer cells to design new drug targets and therapies. A wealth of studies in the past decade have detailed cell autonomous metabolic reprogramming and associated liabilities centering on those processes activated by oncogenes or upon loss of tumor suppressors^2^. More recent studies have built upon this work to describe how the cell of origin influences metabolic programs and liabilities in cancer^3,4^. In addition to these intrinsic programs, properties of the tumor microenvironment can also influence metabolic programs and liabilities in cancer cells^5^. Collectively, these studies have revealed that a common set of genetic alterations can lead to different metabolic dependencies contingent on the tissue type, tumor location, and/or properties of the tumor microenvironment^6–10^.

Previously we found that expression of mutant KRAS, the signature transforming oncogene in pancreatic ductal adenocarcinoma (PDA), rewires central carbon metabolism to support tumor maintenance^11–13^. This includes the diversion of glucose-derived carbon into anabolic pathways that branch from glycolysis and enhanced utilization of glutamine-derived carbon to support anaplerosis in the mitochondria. Of note, these studies demonstrated that oncogenic KRAS enhances activity of the non-oxidative pentose phosphate pathway (PPP), which results in diminished activity of the NADPH-generating oxidative PPP^11^. NADPH is required for the biosynthesis of lipids and deoxynucleotides while simultaneously also serving as an important co-factor to support redox homeostasis. To account for the decreased flux through the oxidative PPP, we reported on a rewired form of the malate-aspartate shuttle that PDA cells utilize to maintain NADPH levels (**Fig. 1a**). This pathway is mediated by the mutant KRAS-driven activation of glutamate oxaloacetate transaminase 1 (GOT1) expression.

**Figure 1:**
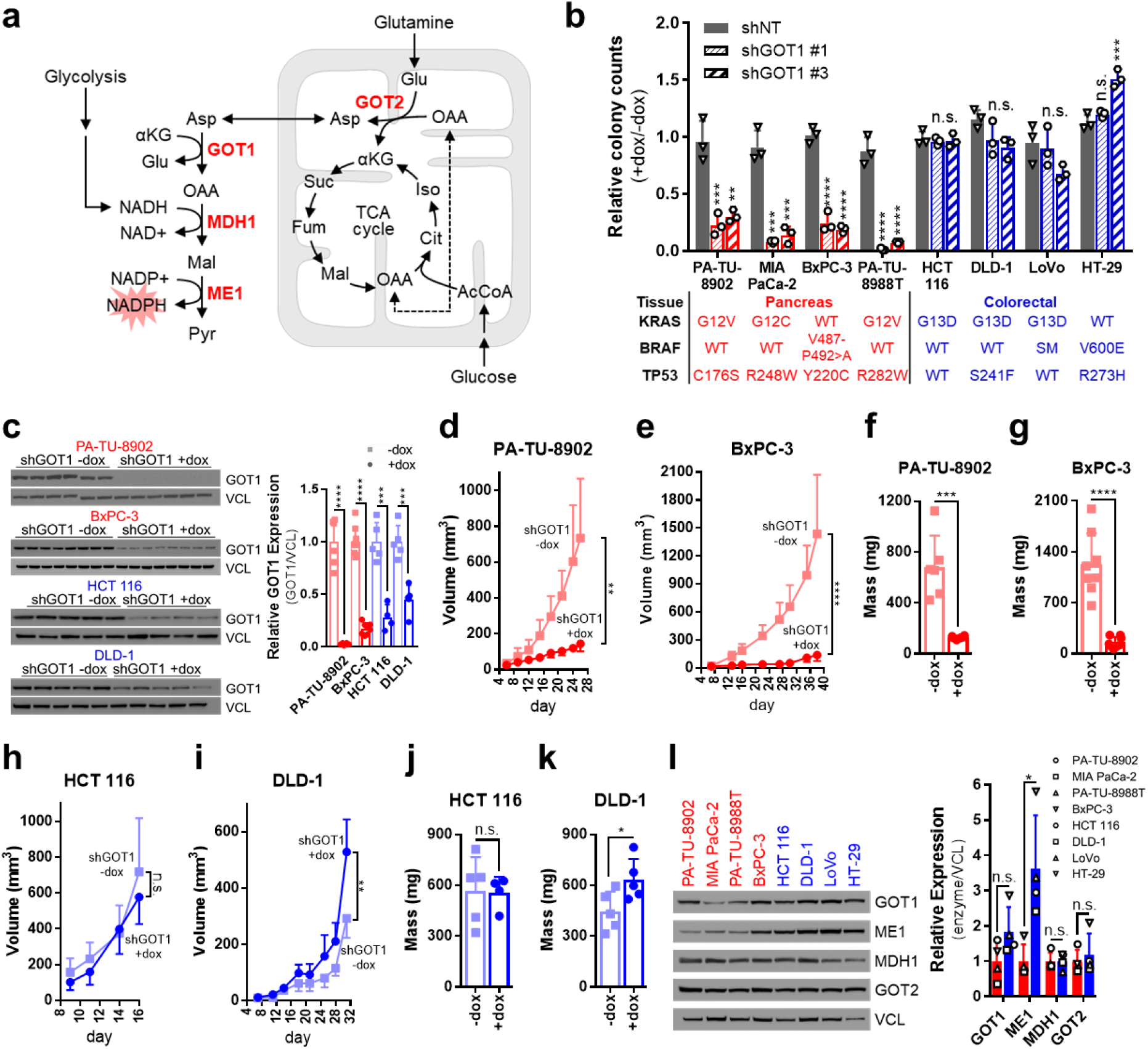
GOT1 dependence exhibits tissue specificity. (**a**) Schematic of the GOT1 pathway in PDA. (**b**) Colony number after dox treatment in PDA (red) and CRC (blue) cell lines expressing dox-inducible (iDox) shRNAs against GOT1 (two independent hairpins; shGOT1 #1, shGOT1 #3) relative to a nontargeting hairpin (shNT). Error bars represent s.d. from biological replicates (n=3). Mutations in *KRAS, BRAF*, and *TP53* are presented in the table below. WT, wildtype; SM, silent mutation. (**c**) Western blots (left) and quantification (right) for GOT1 and vinculin (VCL) loading control from iDox-shGOT1 #1 PDA and CRC tumors. (**d,e**) Tumor growth curves and (**f,g**) final tumor weights from subcutaneous PDA xenografts (n=8, BxPC-3 +/- dox tumors; n=6, PA-TU-8902 +/- dox tumors). Error bars represent s.d. (**h,i**) Tumor growth curves and (**j,k**) final tumor weights from subcutaneous CRC xenografts (n=5, DLD-1 +/- dox, HCT 116 +dox tumors; n=4, HCT 116-dox tumors). Error bars represent s.d. Tumor growth curves for the corresponding iDox-shNT lines are presented in **Extended Fig. 2b**. (**l**) Western blot (left) and quantification (right) for GOT1 pathway components from **Fig. 1a** in wild type PDA and CRC cell lines. AcCoA, acetyl-CoA; αKG, alpha-ketoglutarate; Asp, aspartate; Cit, citrate; Fum, fumarate; Glu, glutamate; GOT1, glutamate oxaloacetate transaminase 1; GOT2, glutamate oxaloacetate transaminase 2; Iso, isocitrate; Mal, malate; MDH1, malate dehydrogenase 1; ME1, malic enzyme 1; NADP+, oxidized nicotinamide adenine dinucleotide phosphate; NADPH, reduced nicotinamide adenine dinucleotide phosphate; OAA, oxaloacetate; Pyr, pyruvate; Suc, succinate. *, *P* < 0.05; **, *P* < 0.01; ***, *P* < 0.001; ****, *P* < 0.0001; Student’s t-test (unpaired, two-tailed).

Importantly, our previous work demonstrated that PDA cells use the NADPH from the GOT1 pathway to manage reactive oxygen species (ROS) through the maintenance of reduced glutathione (GSH) pools^12^. Further, we illustrated that PDA cells were dependent on GOT1 activity for growth in culture, whereas non-transformed fibroblasts and epithelial cells tolerated GOT1 knockdown without consequence. In an effort to leverage these findings about metabolic dependencies in PDA to design new therapies, we recently developed novel small molecule inhibitors that target GOT1^14,15^. Furthermore, GOT1-metabolic pathways have also been shown to play a role in other cancers^16–19^, indicating that GOT1 inhibitors may have utility beyond PDA. However, a rigorous comparison of GOT1 sensitivity in different cancer types has not been performed.

In the current study, we set forth to determine whether the tissue of origin impacts GOT1 dependence to understand which cancers are most likely to benefit from this emerging therapeutic strategy. We found that colorectal cancer (CRC) cell lines harboring *KRAS* and *TP53* mutations, two of the most common mutations in PDA patients^20^, were insensitive to GOT1 inhibition *in vitro* and *in vivo*. This was in dramatic contrast to the PDA models. We then utilized liquid chromatography-coupled mass spectrometry (LC/MS)-based metabolomics strategies, including isotope tracing flux analysis and computational modeling of metabolomics data, to dissect the metabolic consequences of GOT1 knockdown and to contrast how these differed between CRC and PDA cells and tumors. This analysis revealed that GOT1 inhibition uniquely disrupted glycolysis, nucleotide metabolism, and redox homeostasis pathways in PDA. Based on these results, we then designed a combination treatment approach consisting of GOT1 inhibition and radiotherapy. This provided a considerable increase in the efficacy of either single arm treatment uniquely in PDA. Together, these results suggest that the clinical investigation of therapies targeting GOT1, either as monotherapy or in combination with radiation, should begin in PDA. Finally, our data also highlight the importance of tissue of origin in PDA and CRC when studying metabolic wiring and associated dependencies.

## Results

### GOT1 dependence exhibits tissue specificity

To determine whether the tissue of origin impacts GOT1 dependence, we compared GOT1 knockdown in a panel of PDA and CRC cell lines that similarly exhibit mutant KRAS (or BRAF) and mutant TP53 expression (**Fig.1b**). We standardized GOT1 inhibition across experiments by developing doxycycline (dox)-inducible (iDox)-shRNA reagents that target the coding region of GOT1 (shGOT1 #1), the 3’ untranslated region of GOT1 (shGOT1 #3), or a non-targeting shRNA (shNT). shRNA activity was validated after dox administration by assessing GOT1 mRNA and protein expression and intracellular aspartate (Asp), a product of the GOT1 reaction (**Extended Fig. 1a-c**). Additionally, shRNA specificity was validated by rescue with a GOT1 cDNA construct (**Extended Fig. 1d-g**). These constructs were then used to assess GOT1 sensitivity in the panel of PDA and CRC cell lines (**Fig. 1b, Extended Fig. 1h**). As we observed previously with constitutive shRNA targeting GOT1, the colony forming potential of PDA lines was significantly blunted upon inducible GOT1 inhibition. In stark contrast, the CRC cell lines were entirely resistant to growth inhibition in this assay. Importantly, this occurred despite efficient protein knockdown and Asp accumulation in both the PDA and CRC cell lines (**Extended Fig. 1b,c**).

Next, we examined how GOT1 inhibition affected established PDA and CRC tumors. To this end, cells were implanted in the flanks of mice and tumors were allowed to establish for 1 week. Dox was then administered in the chow to initiate GOT1 knockdown (**Fig. 1c**). PDA tumors exhibited a profound retardation of tumor growth (**Fig. 1d-g, Extended Fig. 2a**). Consistent with our *in vitro* observations, CRC lines were insensitive to GOT1 knockdown *in vivo* (**Fig. 1h-k, Extended Fig. 2a**). Xenografts expressing shNT, to control for hairpin and dox effects, showed no difference in growth in either PDA or CRC (**Extended Fig. 2b-e**). The results from these data indicated that, unlike PDA, CRC cell lines and tumors are not dependent on GOT1 for growth.

### Expression of GOT1 pathway components does not distinguish PDA from CRC

Next, we tested if GOT1 dependence was due to lack of, or major differences in, the expression of GOT1-pathway components. Aside from ME1, which showed a modest but statistically significant higher expression in CRC, the GOT1-pathway components examined were expressed at similar levels in the PDA and CRC cells (**Fig. 1l, Extended Fig. 2f**). Importantly, GOT1 is biochemically active in both PDA and CRC cells, as knockdown led to Asp accumulation (**Extended Fig. 1c**). Notably, the Asp build up occurred to a lesser extent in CRC cells compared to PDA cells, suggesting CRC cells may be utilizing compensatory pathways upon GOT1 knockdown. Collectively, these results indicated that while the pathway machinery in PDA and CRC are intact and functional, the differential dependence on GOT1 may result from distinct metabolic pathway activity, rather than enzyme expression, between these two tumor types.

### Differential metabolic pathway activity between PDA and CRC

In order to determine differences in the basal metabolic state between PDA and CRC cells, we used LC/MS-based metabolomics^21–24^ to profile a panel of 3 PDA and 3 CRC parental cell lines in exponential growth phase. Analysis of statistically significant differences in the relative abundance of the steady state metabolite pools indicated that the PDA lines had more gluconodelta lactone-6 phosphate (GdL6P) and 6-phospho gluconate (6PG), metabolites in the oxidative arm of the PPP, and smaller metabolite pools of alanine and lactate (**Fig. 2a**). Many additional differences were observed that did not reach statistical significance, and collectively these revealed an inflection point in glycolysis at the level of aldolase (ALDO).

**Figure 2:**
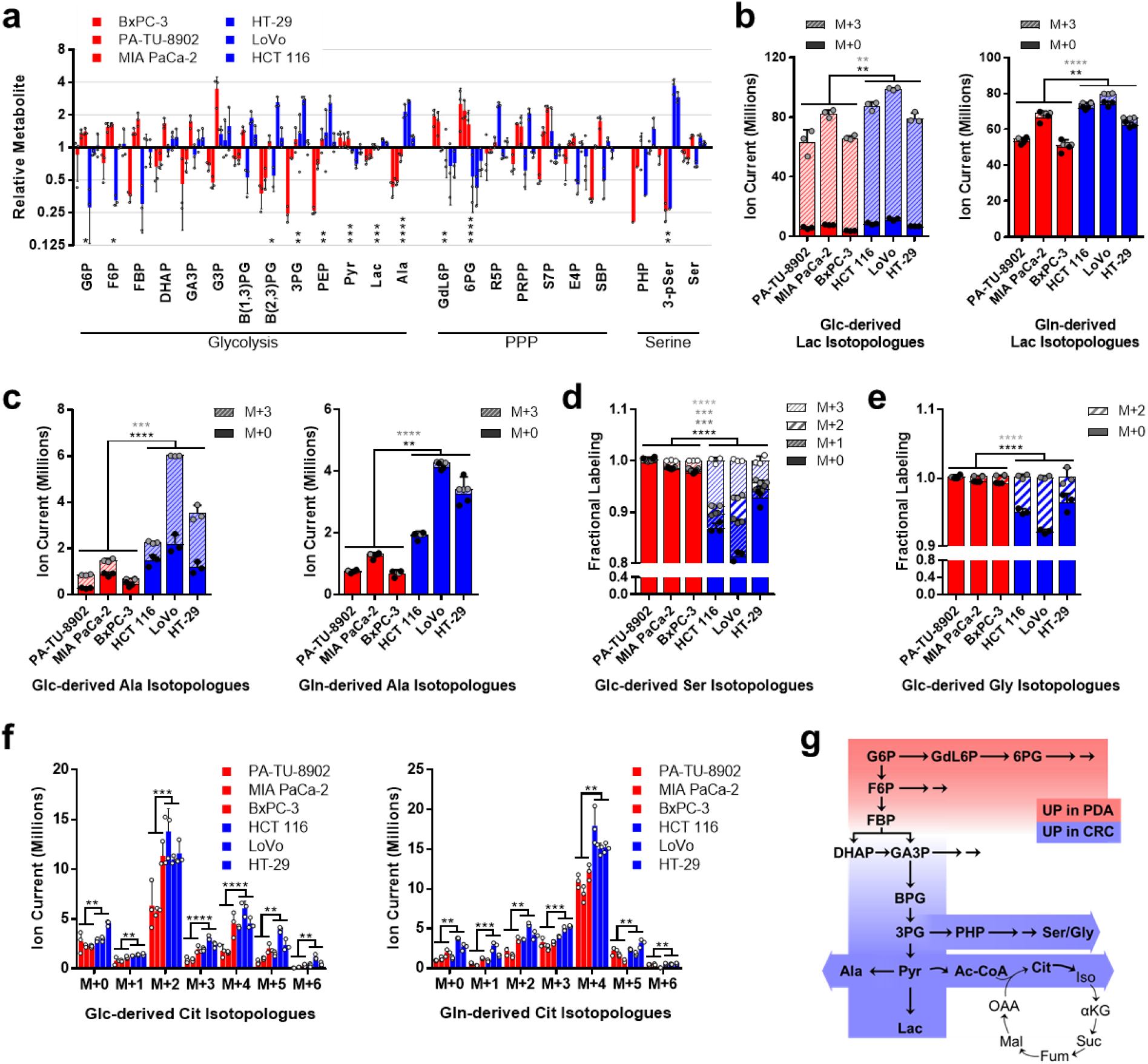
Metabolic profiles of PDA and CRC. (**a**) Relative metabolite levels as determined by LC/MS for glycolysis, pentose phosphate pathway (PPP), and serine metabolism in parental PDA (red) and CRC (blue) cell lines. Uniformly labeled (M+3, hashed bars) and unlabeled (M+0, solid bars) metabolite pools derived from U-^13^C-glucose (Glc, left) or U-^13^C-glutamine (Gln, right) for (**b**) lactate (Lac) and (**c**) alanine (Ala) as determined by LC/MS. (**d**) Relative U-^13^C-Glc-labeling of serine (Ser) and (**e**) glycine (Gly), as determined by gas chromatography (GC)/MS. (**f**) Ion currents for isotopologue distribution of citrate (Cit) derived from U-^13^C-Glc (left) or U-^13^C-Gln (right) in PDA and CRC cell lines. (**g**) Schematic summary of metabolic patterns observed in parental PDA and CRC cells. Red represents increased pool sizes in GOT1-sensitive PDA cells and blue represents increased metabolite pools in GOT1-insensitive CRC cells. 3-pSer, 3-phosphoserine; 3PG, 3-phosphogycerate; 6PG, 6-phosphogluconate; Ac-CoA, acetyl-CoA; αKG, alpha-ketoglutarate; B(1,3)PG, 1,3-bisphosphoglycerate; B(2,3)PG, 2,3-bisphosphoglycerate; Cit, citrate; DHAP, dihydroxyacetone phosphate; E4P, erythrose 4-phosphate; F6P, fructose 6-phosphate; FBP, fructose-1,6-bisphosphate; Fum, fumarate; G3P, glycerol-3-phosphate; G6P, glucose 6-phosphate; GA3P, glyceraldehyde 3-phosphate; GdL6P, glucono-delta-lactone 6-phosphate; Iso, isocitrate; Mal, malate; OAA, oxaloacetate; PEP, phosphoenolpyruvate; PHP, phosphohydroxypyruvate; PRPP, phosphoribosyl pyrophosphate; Pyr, pyruvate; R5P, ribose 5-phosphate; S7P, sedoheptulose-7 phosphate; SBP, sedoheptulose-1,7-bisphosphate; Suc, succinate. Error bars represent s.d. from biological replicates (n=3). Stacked *P*-values are presented for isotopologues in **2b-e** and correspond by color. *, *P* < 0.05; **, *P* < 0.01; ***, *P* < 0.001; ****, *P* < 0.0001; Student t-test (unpaired, two-tailed).

Thus, we set out to further interrogate the metabolic differences between GOT1 dependent and independent cells, and to determine differential central carbon utilization. To this end we performed isotope tracing metabolomics using either uniformly-labeled 13C (U-^13^C) glucose (Glc) or glutamine (Gln)^22–24^ in the parental PDA and CRC lines. Metabolites were collected from log phase cell lines grown overnight in labeled media, and fractional labeling patterns (**Extended Fig. 3**) and metabolite pool sizes (**Extended Fig. 4**) were analyzed.

The fractional labeling patterns between the PDA and CRC cell lines displayed remarkable similarity (**Extended Fig. 3**). In contrast, several notable changes were observed among the relative pool sizes. Similar to our steady state metabolomics (**Fig. 2a**), we observed less lactate (**Fig. 2b**) and alanine (**Fig. 2c**) in the PDA lines, with the majority of this being derived from glucose. Further consistent with the steady state profiling in **Fig. 2a**, the CRC lines have more active serine biosynthetic pathway activity, as illustrated by glucose-derived labeling of serine and glycine (**Fig. 2d,e**). In contrast to these differences, obvious differences in the abundance of Asp, glutamate and alpha-ketoglutarate, the substrates and products of the GOT1 reaction, were not evident between GOT1 dependent and independent lines (**Extended Fig. 4**). This was consistently observed in the glucose and glutamine tracing studies. Similarly, the relative abundance of other TCA cycle intermediates did not exhibit notable differences between the PDA and CRC lines, with the exception of citrate, which is lower in the PDA lines (**Fig. 2f**). These data are summarized together with the unlabeled metabolomic profiling in **Fig. 2g**.

### GOT1 inhibition impairs glycolysis in PDA

It was our expectation that differential GOT1 dependence would be reflected by differences in the baseline wiring of intermediary metabolism between GOT1 dependent and independent parental cell lines. However, given that the steady state profiling data for the unperturbed cells were largely similar (**Extended Figs. 3,4**), we then examined how the metabolome of GOT1 dependent and independent lines responded to knockdown using 3 PDA and 3 CRC iDox-shGOT1 cell lines. In this analysis, we found that Asp increased in all 6 lines and malate decreased in most (**Fig. 3a**), reflecting inhibition of the GOT1 pathway^12^. In addition, all 6 lines showed a consistent accumulation of glycolytic intermediates between the ALDO-catalyzed and pyruvate kinase (PK)-catalyzed steps of glycolysis (**Fig. 3b,c**). Despite these consistencies, GOT1 knockdown only impaired glycolytic flux in PDA, as measured by Seahorse Metabolic Flux Assay (**Fig. 3d**).

**Figure 3:**
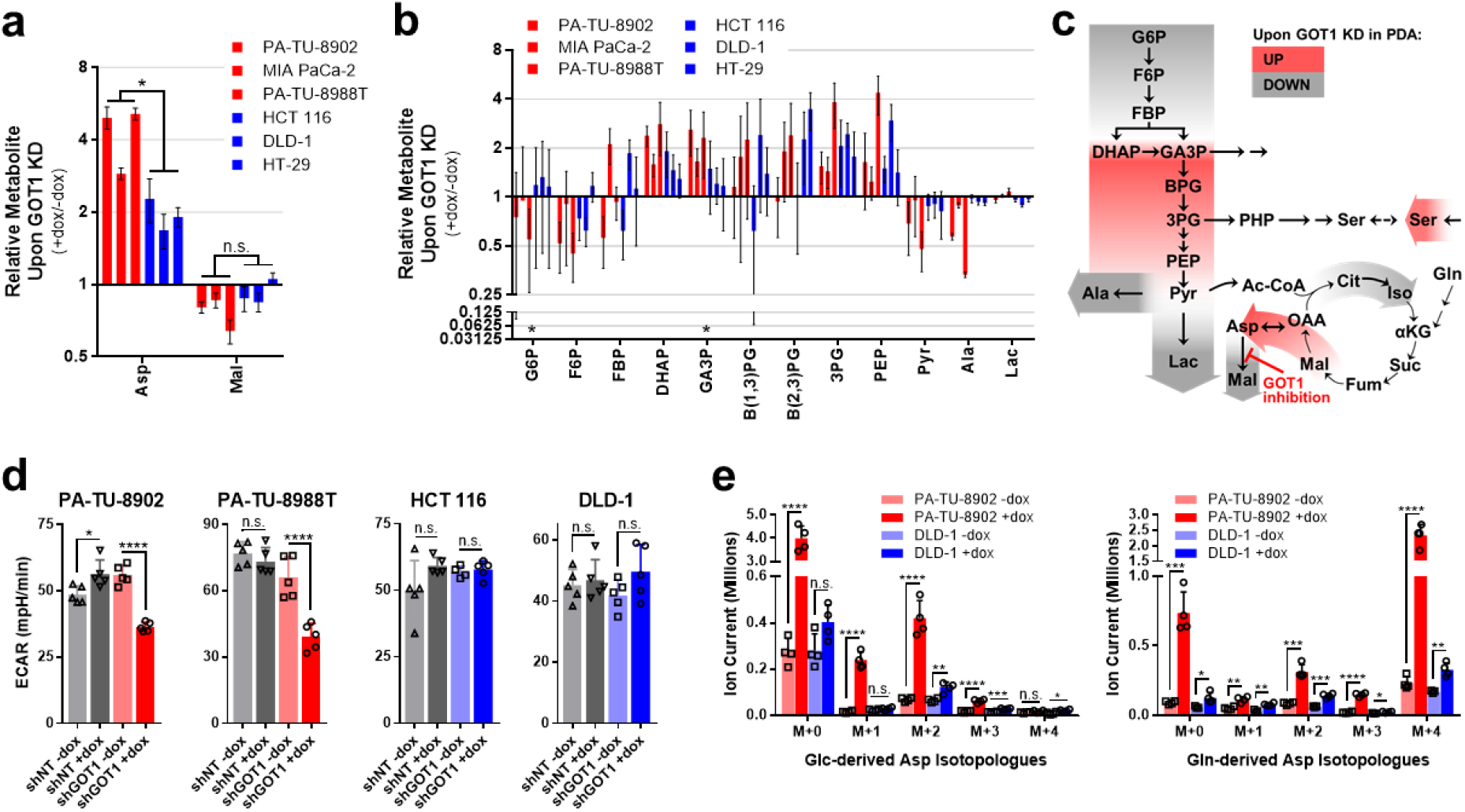
Metabolic profiles of PDA and CRC following GOT1 inhibition. (**a**) Relative aspartate (Asp) and malate (Mal) pools as determined by LC/MS in iDox-shGOT1 #1 PDA and CRC, presented as GOT1 knockdown over mock (+dox/-dox). (**b**) Relative glycolysis metabolite pools, as presented in **3a**. (**c**) Summary of changes to central carbon metabolism upon GOT1 knockdown PDA cells. Red represents increased pool sizes in PDA cells upon GOT1 knockdown and green represents decreased metabolite pools. (**d**) Basal extracellular acidification rate (ECAR) levels in iDox-shGOT1 #1 and control shNT PDA and CRC cells, as determined by Seahorse Metabolic Flux Analysis. (**e**) Ion currents for isotopologue distribution of Asp derived from U-^13^C-Glc (left) or U-^13^C-Gln (right) in iDox-shGOT1 #1 PA-TU-8902 PDA and DLD-1 CRC cell lines. 3PG, 3-phosphogycerate; Ac-CoA, acetyl-CoA; αKG, alpha-ketoglutarate; Ala, alanine; B(1,3)PG, 1,3-bisphosphoglycerate; B(2,3)PG, 2,3-bisphosphoglycerate; Cit, citrate; DHAP, dihydroxyacetone phosphate; F6P, fructose 6-phosphate; FBP, fructose-1,6-bisphosphate; Fum, fumarate; G6P, glucose 6-phosphate; GA3P, glyceraldehyde 3-phosphate; Iso, isocitrate; Lac, lactate; OAA, oxaloacetate; PEP, phosphoenolpyruvate; Pyr, pyruvate; Suc, succinate. Error bars represent s.d. from biological replicates (n=3, in **a**, **b**, **e**; n=5 in **d**). *, *P* < 0.05; **, *P* < 0.01; ***, *P* < 0.001; ****, *P* < 0.0001; Student t-test (unpaired, two-tailed).

To further interrogate these metabolic differences, we also employed ^13^C-Glc and Gln tracing analyses following GOT1 knockdown in the PA-TU-8902 PDA and DLD-1 CRC lines (**Extended Figs. 5,6**). In cells of both tissue types, glycolytic intermediates were entirely Glc-derived, TCA cycle intermediates were predominantly Gln-derived, and GOT1 knockdown did not promote differential nutrient utilization to fuel these pathways (**Extended Figs. 5,6**). As expected, pronounced accumulation of Asp was observed, and, as we have seen previously^12^, this is predominantly derived from Gln in cultured cells (**Fig. 3e**). Again, the fractional labeling data indicate largely consistent patterns of metabolite changes and nutrient utilization in glycolysis and the TCA cycle, and yet despite this, glycolytic activity and proliferation are only impaired in the PDA cells (**Fig. 1b, 3d**).

### GOT1 inhibition disrupts nucleotide metabolism in PDA cells

The growth inhibitory activity of GOT1 knockdown in PDA has prompted ongoing efforts to develop small molecule GOT1 inhibitors^14,15^. To further harness the GOT1 selective dependence of PDA, we sought to identify metabolic pathways that could be targeted in combination with GOT1. Thus, to look more broadly at how GOT1 knockdown impacts metabolism between GOT1-dependent PDA and -independent CRC cell lines, we analyzed the unlabeled metabolomics data, as follows. The ~250 metabolites across central carbon metabolism were plotted as the average of the 3 PDA lines (dox/mock) over the average of the 3 CRC lines (dox/mock) (**Fig. 4a**). We identified pathways that are uniquely disrupted upon GOT1 knockdown in the PDA lines by analyzing metabolites with a greater than 2-fold change via MetaboAnalyst Pathway Analysis^25^. Among the differentially represented pathways, we observed that pyrimidine and purine metabolism were the most significantly enriched between PDA and CRC cell lines (**Extended Fig. 7a**). Metabolites from PDA and CRC xenografts were analyzed in a similar manner with pyrimidine and purine metabolism also significantly enriched (**Extended Fig. 7b**). We also found that several nodes in nucleotide metabolism were deregulated in PDA cells upon GOT1 inhibition by modeling our metabolomics data with the Recon1 genome-scale network model^26,27^ with dynamic flux analysis (DFA)^28,29^ (**Extended Fig. 7c,d, Extended Table 1**). Given the importance of nucleic acid metabolism in proliferation and the response to damage, we hypothesized that GOT1 inhibition would modulate the cellular response to additional perturbations to these pathways.

**Figure 4:**
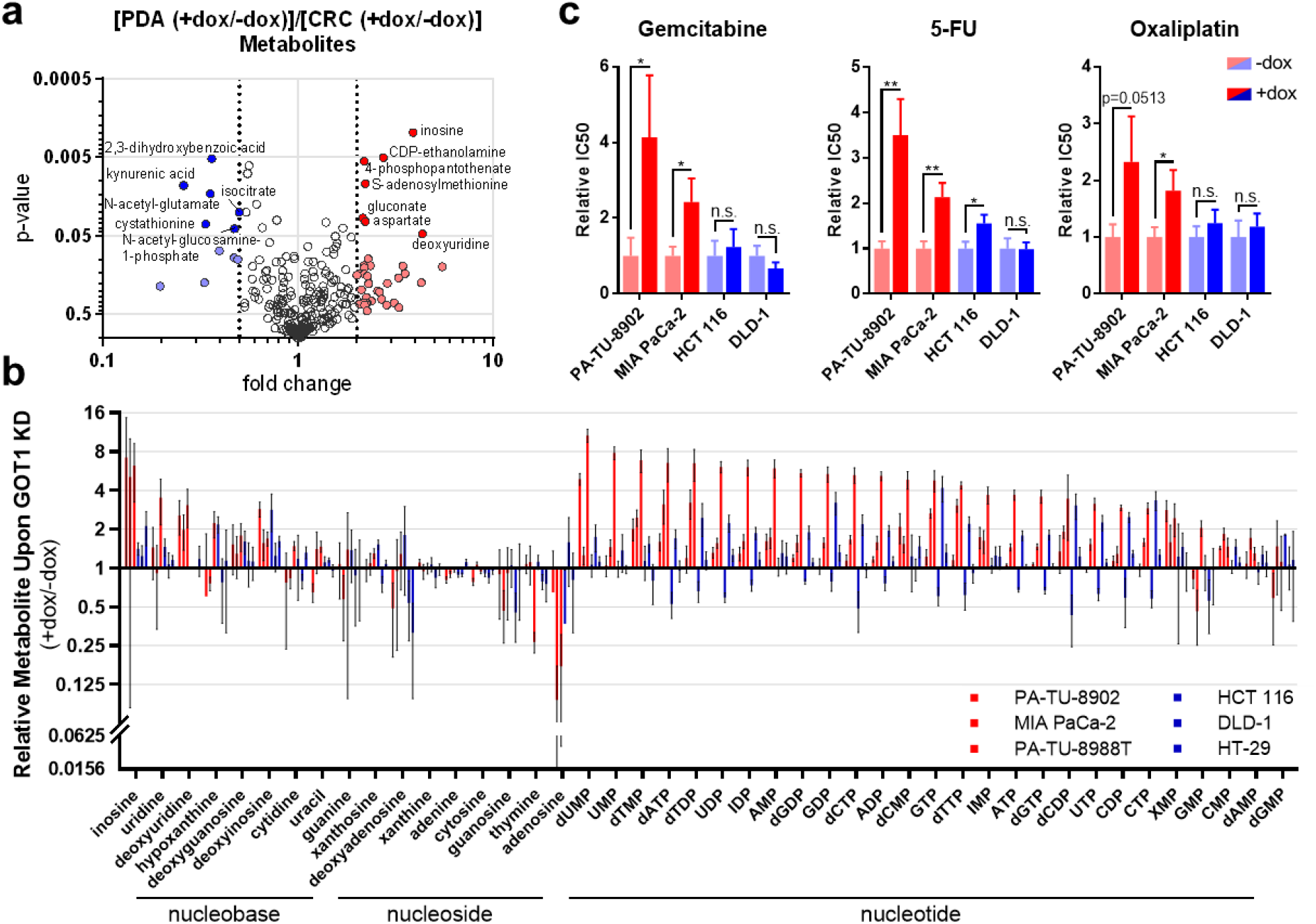
GOT1 inhibition disrupts nucleotide metabolism in PDA. (**a**) Fold-change versus p-value are plotted per metabolite as the average from 3 iDox-shGOT1 #1 PDA lines (+dox/-dox) over the average from 3 iDox-shGOT1 #1 CRC lines (+dox/-dox). Metabolites with filled circles were used for the pathway analysis in **Extended Fig. 6a**. Metabolites identity is indicated for those with *P* < 0.05 and fold change +/- 2. (**b**) Relative nucleic acid pools as determined by LC/MS in iDox-shGOT1 #1 PDA and CRC, presented as GOT1 knockdown over mock (+dox/-dox). (**c**) Relative IC50 of gemcitabine, 5-fluorouracil (5-FU) and oxaliplatin in iDox-shGOT1 #1 PDA and CRC cells upon GOT1 knockdown. The dose response curves from which the IC50s were derived are presented in **Extended Fig. 7e**. ADP, adenosine diphosphate; AMP, adenosine monophosphate; ATP, adenosine triphosphate; CDP, cytidine diphosphate; CMP, cytidine monophosphate; CTP, cytidine triphosphate; dAMP, deoxyadenosine monophosphate; dATP, deoxyadenosine triphosphate; dCDP, deoxycytidine diphosphate; dCMP, deoxycytidine monophosphate; dCTP, deoxycytidine triphosphate; dGDP, deoxyguanosine diphosphate; dGMP, deoxyguanosine monophosphate; dGTP, deoxyguanosine triphosphate; dTDP, deoxythymidine diphosphate; dTMP, deoxythymidine monophosphate; dTTP, deoxythymidine triphosphate; dUMP, deoxyuridine monophosphate; GDP, guanosine diphosphate; GMP, guanosine monophosphate; GTP, guanosine triphosphate; IDP, inosine diphosphate; IMP, inosine monophosphate; UDP, uridine diphosphate; UMP, uridine monophosphate; UTP, uridine triphosphate; XMP, xanthosine monophosphate. Error bars in **b**,**c** represent s.d. from biological replicates (n=3). n.s., not significant; *, *P* < 0.05; **, *P* < 0.01; Student t-test (unpaired, two-tailed).

### GOT1 inhibition protects PDA cells from cytotoxic chemotherapy

Gemcitabine and 5-fluorouracil (5-FU) are pyrimidine analogs and front-line chemotherapies used to treat PDA patients^30–32^. Inspection of pyrimidine metabolism in our datasets revealed that it scored among the top differentially active pathways in both the MetaboAnalyst and DFA. Accordingly, we analyzed the unlabeled metabolomics data for nucleobase, nucleoside, and nucleotide pool levels after GOT1 knockdown and found that many are increased in PDA cellscompared to CRC cells (**Fig. 4b**). We hypothesized that the increase in these metabolites upon GOT1 inhibition may serve to compete with anti-metabolite based therapies, as we have seen in other contexts^24,33,34^. To test this hypothesis, we treated PDA and CRC lines with a dose response of gemcitabine and 5-FU-in the presence or absence of GOT1 inhibition. We also included oxaliplatin, a mechanistically distinct alkylating agent used in PDA front-line therapy. In line with our hypothesis, GOT1 knockdown in PDA cells promoted resistance to gemcitabine and 5-FU, whereas knockdown did not similarly impact resistance to chemotherapy in the CRC lines (**Fig. 4c, Extended Fig. 7e**).

### GOT1 inhibition decreases GSH and sensitizes PDA cells to radiation therapy

Cysteine and sulfur metabolism were the next most deregulated pathways between GOT1 inhibited PDA and CRC cells (**Extended Fig. 7a**). In tumors, cysteine metabolism was the third most significantly enriched pathway (**Extended Fig. 7b**) and DFA shows GSH metabolism is a vulnerability in PDA (**Extended Fig 7c, d**). Cysteine is the rate limiting amino acid in GSH biosynthesis, and in our previous studies, we observed a drop in GSH pools following GOT1 knockdown^12^. Thus, we directed our attention to changes in GSH between PDA and CRC lines. Here, as determined by LC/MS, we found that both GSH and the reduced to oxidized glutathione (GSSG) ratio (GSH/GSSG), were uniquely decreased in PDA cells (**Fig. 5a**). The decrease in the GSH/GSSG ratio was similarly observed using a biochemical assay in our panel of 3 PDA and 3 CRC cell lines (**Fig. 5b**). Furthermore, we also observed that the GSH/GSSG ratio decreased as a function of the duration of GOT1 knockdown, which similarly paralleled with increasing levels of Asp (**Fig. 5c-e, Extended Fig. 8a,b**).

**Figure 5:**
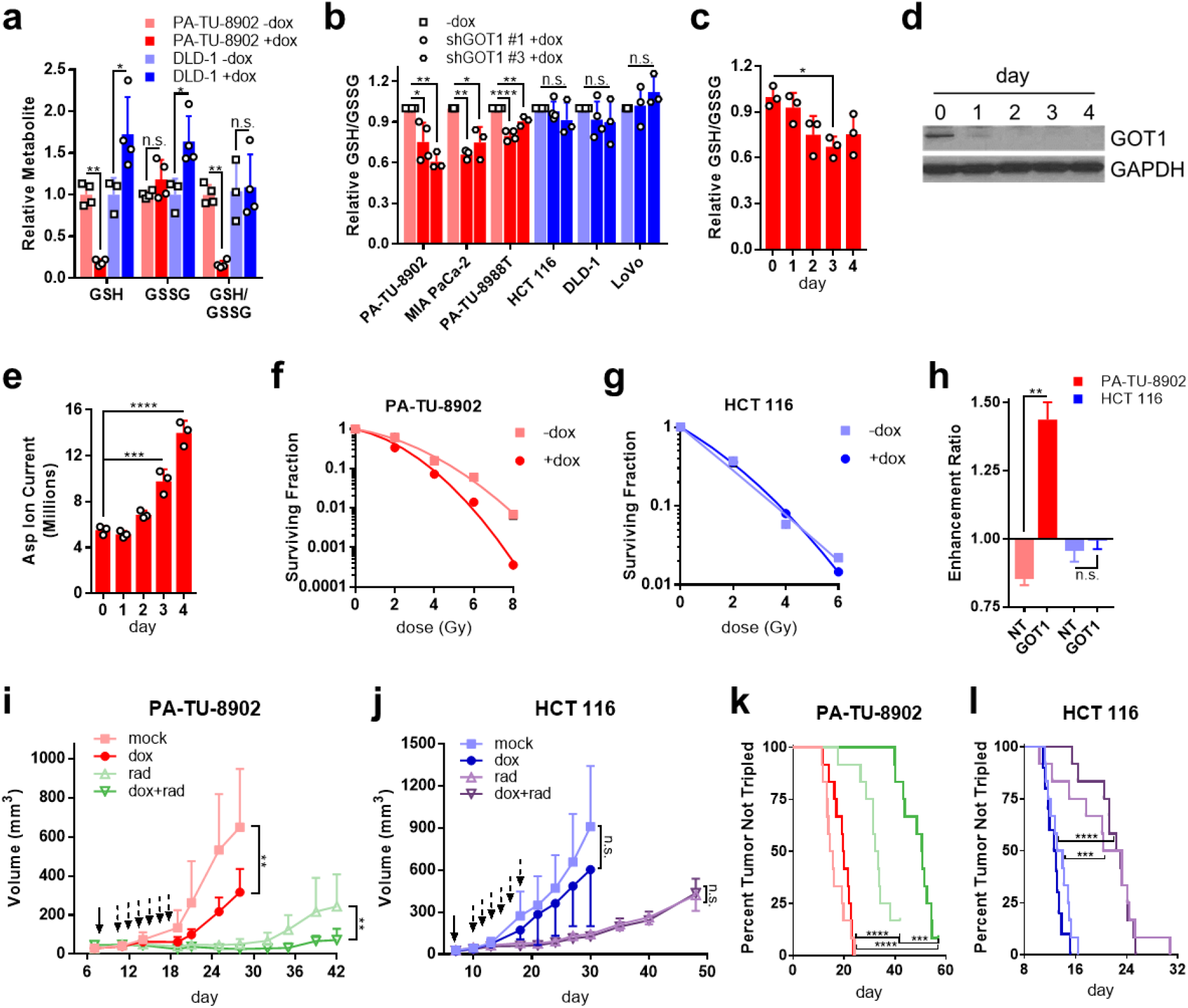
GOT1 inhibition induces redox imbalance and sensitizes PDA to radiation therapy. (**a**) Relative reduced glutathione (GSH), oxidized glutathione (GSSG), and the ratio (GSH/GSSG) pools upon GOT1 knockdown in iDox-shGOT1 #1 PDA and CRC cells. Data were obtained by LC/MS and are normalized as GOT1 knockdown over mock (+dox/-dox). (**b**) Relative GSH/GSSG upon GOT1 knockdown as determined by enzymatic assay in PDA and CRC and normalized as GOT1 knockdown over mock (+dox/-dox). (**c**) Timecourse of relative GSH/GSSG pools upon GOT1 knockdown in iDox-shGOT1 #1 PA-TU-8902 PDA cells. Data were obtained by LC/MS and are normalized as GOT1 knockdown over mock (+dox/-dox). (**d**) GOT1 protein expression during dox-mediated knockdown timecourse in iDox-shGOT1 #1 PA-TU-8902 cells. GAPDH serves as the protein loading control. (**e**) Total aspartate (Asp) pools in PA-TU-8902 cells, as determined by LC/MS and plotted as GOT1 knockdown over mock (+dox/-dox). (**f**) Surviving fraction from clonogenic cell survival assays of radiation-treated iDox-shGOT1 #1 PA-TU-8902 and (**g**) HCT 116. Gy, Gray. (**h**) Enhancement ratio of radiation treated iDox-shGOT1 #1 PDA and CRC cells. Error bars represent s.d. from biological replicates in **a,c-h** (n=3) and in **b** (n=4, iDox-shGOT1 #1 PA-TU-8902, PA-TU-8988T, HCT 116, DLD-1; n=3, iDox-shGOT1 #1 MIA PaCa-2, LoVo and iDox-shGOT1 #3). (**i**) Tumor growth of iDox-shGOT1 #1 PA-TU-8902 or (**j**) iDox-shGOT1 #1 HCT116 xenografts treated with dox (solid arrow; maintained for the duration of the experiment) and/or radiation (rad; dashed arrows). n=12 tumors per arm, except for dox HCT 116 tumors where n=10. Error bars represent s.d. (**k**) Time to tumor tripling of iDox-shGOT1 #1 PA-TU-8902 or (**l**) iDox-shGOT1 #1 HCT116 xenografts. n.s., not significant; *, *P* < 0.05; **, *P* < 0.01; ***, *P* < 0.001; ****, *P* < 0.0001; Student t-test (unpaired, two-tailed) (**a,h,I,j**); one-way ANOVA (**b,c,e,k,l**).

Radiotherapy is a pro-oxidant treatment modality frequently used to treat locally advanced PDA, but its efficacy can be limited by both the intrinsic treatment resistance of PDA and the risk of inducing toxicity in the nearby small bowel^35,36^. Given that radiation induces cell death through oxidative damage to DNA, and its effects can be mitigated by high levels of antioxidants such as GSH, we hypothesized that GOT1 inhibition would selectively radiosensitize PDA with minimal effects in other tissues that do not depend on GOT1 to maintain redox balance. To test this, we first examined the response of PDA to radiation using an *in vitro* clonogenic assay (**Fig 5f**). This demonstrated that PDA cells were sensitized to radiation after dox-induced GOT1 knockdown, whereas no effect was observed in CRC cells (**Fig. 5g,h**). Importantly, this effect was not observed in controls (**Extended Fig. 8c**). GOT1 knockdown provided a radiation enhancement ratio of 1.4, a similar score observed with classical radiosensitizers^37–39^ (**Fig 5h**).

Based on these results, we then explored the utility of GOT1 inhibition as a radiosensitizing strategy in PDA and CRC tumor models *in vivo*. PDA or CRC tumors were established as in **Fig. 1d-k**, with radiation treatment administered in 6 daily doses beginning on day 10. GOT1 knockdown significantly impaired tumor growth in PDA but not CRC (**Fig. 5i,j, Extended Fig. 8d,e**). Radiotherapy was efficacious as a single agent in both models and delayed tumor growth. However, GOT1 inhibition uniquely increased the time to tumor tripling in PDA (**Fig. 5k,l**). Together with our mechanistic studies above, these results demonstrate that GOT1 inhibition promotes redox imbalance uniquely in PDA, which results in a drop in GSH levels and the GSH/GSSG ratio, leading to radiosensitization of PDA cells *in vitro* and PDA tumors *in vivo*.

## Discussion

Precision oncology aims to assign new medicines based on the genotype of a patient^40^. In PDA and CRC, activating mutations of the MAPK pathway (e.g. in *KRAS* and *BRAF)* and loss of tumor suppressor *TP53* are common^41,42^, and these mutations play important roles in the reprogramming of cancer metabolism^11,18,43^. Yet, despite this, metabolic gene expression programs in tumors more closely resemble their cell of origin rather than their oncogenotype^44^. Our results similarly add to the growing body of literature that metabolic dependencies exhibit tissue specificity^3,4^. Herein, we report that among typically mutant KRAS-expressing PDA and CRC lines, PDA cells are uniquely responsive to GOT1 knockdown. This is manifest as profound growth inhibition *in vitro* and in tumor xenografts *in vivo*. Through an integrated analysis utilizing multiple metabolomics profiling approaches together with computational modeling, we demonstrate that GOT1 knockdown uniquely impacts glycolysis, nucleotide metabolism, and GSH-mediated redox regulation in PDA. Based on the disrupted GSH profile, we demonstrated that GOT1 knockdown can serve as a radiosensitizing strategy for PDA.

Despite observing a stark difference in the GOT1 dependence between our PDA and CRC cell lines, the baseline metabolic profiles and nutrient utilization in central carbon metabolism was surprisingly comparable (**Fig. 2, Extended Figs. 3,4**). This similarity may reflect adaptations that have occurred during prolonged exposure of these cell lines to culture, which serve to meet an optimal metabolic flux program to facilitate maximal proliferation. Regardless, metabolic dependences remain hard-wired. Upon GOT1 inhibition, unique shifts in metabolism are observed between the two tissue types, which account for the growth inhibition of PDA cells and tumors upon GOT1 knockdown.

A disruption of nucleotide metabolism was the most notable metabolic change across PDA lines upon GOT1 knockdown (**Fig. 4b, Extended Fig. 7a**). Generally, this led to the accumulation of numerous phosphorylated nucleic acid species. We also observed that GOT1 knockdown reduced the sensitivity of PDA cells to the anti-metabolite chemotherapies gemcitabine and 5-FU. Our proposed explanation for these results is that the increase in the pool of deoxycytidine and uracil species, respectively, decreases the relative intracellular concentration of the anti-metabolite therapies and thereby their activity^24,33^. This explanation, however, does not apply to oxaliplatin, whose cytotoxic activity is similarly impaired upon GOT1 knockdown in PDA cells. Thus, a non-mutually exclusive explanation for the chemoprotective effect of GOT1 is that GOT1 knockdown is cytostatic in PDA cells and tumors (**Fig. 1b,c**). Chemotherapy is thought to work by selectively targeting dividing cells. Given that GOT1 knockdown impairs proliferation, the chemoprotective effect may simply result from impairing cycling, an effect not observed in GOT1 inhibition resistant CRC. Collectively, these results indicated that the use of chemotherapy in conjunction with GOT1 is not a practical therapeutic strategy and highlights the need to test combination treatment strategies in the preclinical setting.

PDA is an extremely aggressive disease and therapeutic options are largely ineffective^45^. The odds of surviving the first year are only 24%, and the five-year survival rate is a dismal 9%^46^. One of the main factors underscoring this low survival rate is the lack of effective clinical treatments^47^. *KRAS* mutations are observed in >90% of PDA, yet despite great efforts, current means to inhibit RAS are limited to the G12C mutation^48^, which is only observed in 2% of patients^41^. Immunotherapy, while promising in other types of cancer, has proven ineffective to treat PDA^49,50^. Thus, improving current therapeutic modalities represents the best immediate hope for PDA patients. Radiotherapy is a standard of care for PDA in many institutions, although this remains controversial^51^. For patients that have undergone surgical resection for PDA, the receipt of adjuvant radiation (in combination with chemotherapy) is associated with a survival benefit in large institutional series, and this is currently being evaluated in a phase III randomized trial (RTOG 0848, ClinicalTrials.gov NCT01013649)^52,53^. Despite these encouraging results, nearly 40% of patients receiving adjuvant radiation experience treatment failures within the irradiated field, indicating that PDA radiation resistance remains an important barrier to improving outcomes in the adjuvant setting^54^. Radiation also plays an important role for patients with locally advanced PDA that cannot be resected^55,56^. As in the adjuvant setting, nearly 40% of patients receiving radiation for locally advanced PDA will experience local tumor progression, again highlighting the clinical challenge of radiation resistance in PDA^57^.

Our findings suggest that GOT1 inhibition could improve outcomes in PDA by overcoming this radiation resistance. Importantly, this strategy is unlikely to increase the normal tissue toxicity that often limits the intensification of radiotherapy-based treatment regimens. This potential therapeutic window is supported by our previous reports that GOT1 inhibition is well tolerated in non-transformed cells, as radiation dose is often limited so as not to harm nearby normal tissues. The GOT1 independence of CRC cell lines provide further support that a therapeutic window may exist for systemically targeting GOT1 in a subset of cancer types. To this end, we and others have engaged in developing GOT1 inhibitors^14,15^. Future work on optimized GOT1 drugs will be required to test the activity of these agents in combination with radiotherapy.

## Supporting information

Nelson, et al_Extended Figures

## Acknowledgements

BSN was supported by T32CA009676 and T32DK094775; DMK by the University of Michigan’s Program in Chemical Biology Graduate Assistance in Areas of National Need (GAANN) award; ZCN by U066362; CJH by P30DK034933, UL1TR000433, T32CA009676, and F32CA228328; JMA and LCC by NIH grant 2P01CA120964; LCC by grant R35-CA197588; HCC by the Sky Foundation; CAL, HCC, and DRW by a Cancer Center Support Grant (P30CA046592); CAL and HCC by U01 CA224145; CAL by the Pancreatic Cancer Action Network/AACR (13-70-25-LYSS), Damon Runyon Cancer Research Foundation (DFS-09-14), V Foundation for Cancer Research (V2016-009), Sidney Kimmel Foundation for Cancer Research (SKF-16-005), the AACR (17-20-01-LYSS), and an ACS Research Scholar Grant (RSG-18-186-01). Metabolomics studies at U-M were supported by DK097153.

## Author Contributions

BSN, LL, LCC, AK, DMW, and CAL conceived of and designed this study. BSN and CAL planned and guided the research and wrote the manuscript. BSN, LL, DMK, CS, CCR, AM, JR, TG, IK, KWR, JD, MD, ZN, OM, BR, ZT, CH, JA, HC, AK, DMW, and CAL performed experiments, analyzed, and interpreted data. AK, DMW, and CAL supervised the work carried out in this study.

## Materials and Methods

### Cell culture

Cell lines were obtained from the American Type Culture Collection or the German Collection of Microorganisms and Cell Cultures: PDA cell lines PA-TU-8902 (RRID:CVCL_1845), BxPC-3 (RRID:CVCL_0186), MIA PaCa-2 (RRID:CVCL_0428), and PA-TU-8988T (RRID:CVCL_1847); and CRC cell lines HCT 116 (RRID:CVCL_0291), DLD-1 (RRID:CVCL_0248), LoVo (RRID:CVCL_0399), and HT-29 (RRID:CVCL_0320). All cell lines were routinely tested for mycoplasma contamination (Lonza MycoAlert Plus, LT07-710). BxPC-3 cells were cultured in RPMI-1640 (Gibco, 11875-093) with 10% FBS (Corning, 35-010-CV). All other cell lines were cultured in DMEM (Gibco, 11965-092) with 10% FBS.

### shRNA constructs and iDox-shRNA stable cell lines

The lentiviral vector containing tetracycline inducible system Tet-pLKO-puro (a gift from Dmitri Wiederschain) was engineered to contain the following shRNAs: GOT1 coding region (shGOT1 #1, TRCN0000034784) or GOT1 3’UTR (shGOT1 #3, 5’-CCGGTTGGAGGTCAAAGCA AATTAACTCGAGTTAATTTGCTTTGACCTCCAATTTTT-3’). Oligonucleotides were obtained (Integrated DNA Technologies Inc.), annealed and cloned at AgeI and EcoRI sites in tet-pLKO-puro (Addgene, 21915; http://www.addgene.org/21915, RRID:Addgene_21915)^58^ following the Wiederschain Protocol (https://media.addgene.org/data/plasmids/21/21915/21915-attachment_Jws3xzJOO5Cu.pdf). A tet-pLKO non-targeting control vector (shNT, 5’-CCGGCAACAAGATGAAGAGCACCAACTCG AGTTGGTGCTCTTCATCTTGTTGTTTTT-3’; or shLUC, TRCN0000072259) was constructed similarly. Tet-pLKO-shGOT1 and tet-pLKO-shNT lentiviruses were produced by the University of Michigan Vector Core using the purified plasmids. Parental PDA and CRC cell lines were then transduced with optimized viral titers and stable cell lines were established post puromycin selection.

### Colony forming and clonogenic cell survival assays

Colony forming assays (CFA) were performed as previously described with slight modifications^12^. Briefly, cells were plated in 6-well plates at 300-600 cells per well (dependent on the cell line) in 2 mL of media. 24 hours after seeding, dox was added at 1 ug/mL and culture medium was changed every 48 hours. After 8-13 days, colonies were fixed with 100% methanol and stained with 0.5% crystal violet solution. Colonies in triplicate wells were counted in ImageJ and graphed. Statistical analyses performed using GraphPad Prism7 software.

For radiotherapy studies, clonogenic assays were performed as described previously^59,60^. Briefly, 3 to 4 days after dox-induced shRNA expression, cells were irradiated with varying doses of radiation and then replated at clonal density. After 10 to 14 days of growth, colonies of 50 or more cells were enumerated and corrected for plating efficiency using unirradiated samples. Cell survival curves were fitted using the linear-quadratic equation. Enhancement ratios were calculated as the ratio of the mean inactivation dose under no dox conditions divided by the mean inactivation dose under +dox conditions.

### cDNA rescues

Direct mutagenesis of shGOT1 #1 in pDONR223 resulted in GOT1 cDNA (sequence GCGGTGGTATAACGGCACCAA) resistant to shRNA targeting. GOT1 cDNA was Gateway cloned into DEST vector pLVX-GW-Hygro.

### qPCR

RNA was extracted using RNeasy Mini Kit (Qiagen, 74104) according to manufacturer’s instructions. cDNA was generated using SuperScript III CellsDirect™ cDNA Synthesis Kit (Invitrogen, 18080300). RT–PCR was done using SYBR Green PCR Master Mix (Applied Biosystems, 4309155) on a ViiA 7 Real-Time PCR System (Applied Biosystems). Relative mRNA levels were normalized to expression of human β-actin. RT–PCR was performed in quadruplicate.

### Western blot analysis

Stable shNT and shGOT1 cells were cultured with or without dox media and protein lysates were collected after five days using RIPA buffer (Sigma, R0278) containing protease inhibitor cocktail (Sigma/Roche, 04 693 132 001). Samples were quantified with Pierce BCA Protein Assay Kit (ThermoFisher, 23225). 10 to 40 μg of protein per sample were resolved on NuPAGE Bis-Tris Gels (Invitrogen, NP0336) and blotted to PVDF membranes (Millipore, IPVH00010). Membranes were blocked in Tris-buffered saline (Bio-Rad, 170-6435) containing 0.5% of Tween 20 (Sigma, P2287) (TBS-T buffer) and 5% non-fat dry milk (LabScientific, M0841) then incubated with primary antibody overnight at 4°C. The membranes were then washed with TBS-T buffer followed by exposure to the appropriate horseradish peroxidase-conjugated secondary antibody for 1h and visualized on either Kodak X-ray film (GeneMate, F-9023-8×10) or BioRad ChemiDoc Imaging System using either SuperSignal West Pico Chemiluminescent Substrate (Thermo Scientific, 34080) or ECL Prime Western Blotting Detection Reagent (Amersham, RPN2232). The following antibodies were used: anti-aspartate aminotransferase (anti-GOT1) at a 1:1,000 dilution (Abcam, ab171939), anti-GOT2 at a 1:1,000 dilution (Atlas Antibodies Sigma-Aldrich, HPA018139), anti-ME1 at a 1:1,000 dilution (Santa Cruz, sc-100569), anti-MDH1 at a 1:10,000 dilution (Abcam, ab180152), and loading control vinculin at a 1:10,000 dilution (Cell Signaling Technology, 13901) or GAPDH at a 1:1,000 dilution (Cell Signaling Technology, 2118). Anti-GOT1 at a dilution of 1:1,000 (Abnova, H00002805-B01P) was used in **Fig. 5d** and **Extended Fig. 1d**. Anti-rabbit IgG, HRP-linked (Cell Signaling Technology, 7074) and anti-mouse IgG, HRP-linked (Cell Signaling Technology, 7076) secondary antibody was used at a 1:10,000 dilution. Protein expression was quantified with ImageJ.

### Mass Spectrometry-Based Metabolomics

#### Unlabeled targeted metabolomics

Cells were plated at 500,000 cells per well in 6-well plates or ~1.5 million cells per 10 cm dish. At the end of indicated time points, 1 mL of medium was saved for metabolite extraction. Cells were lysed with dry-ice cold 80% methanol and extracts were then centrifuged at 10,000 g for 10 min at 4°C and the supernatant was stored at −80°C until further analyses. Protein concentration was determined by processing a parallel well/dish for each sample and used to normalize metabolite fractions across samples. Based on protein concentrations, aliquots of the supernatants were transferred to a fresh micro centrifuge tube and lyophilized using a SpeedVac concentrator. Dried metabolite pellets from cells or media were re-suspended in 35 μl 50:50 methanol:water mixture for LC–MS analysis. Data was collected using previously published parameters (ref:^22^).

Raw data were pre-processed with Agilent MassHunter Workstation Software Quantitative QqQ Analysis Software (B.07.00). Additional statistical analyses were carried out in Excel (Microsoft) where each sample was normalized by the total intensity of all metabolites to reflect the protein content as a normalization factor. We then retained only those metabolites with at least 2 replicate measurements. The remaining missing value in each condition for each metabolite was filled with the median value of the other replicate measurements. Finally, each metabolite abundance level in each sample was divided by the median of all abundance levels across all samples to obtain relative metabolites. Significance testing was a two-tailed t-test with a significance threshold level of 0.05.

#### 13C-tracing analysis

Cells were cultured in DMEM lacking Glc or Gln (ThermoScientific, A1443001) and supplemented with 10% dialyzed FBS (ThermoScientific, 26400036), the appropriate labeled substrate U-^13^C-Gln (Cambridge Isotope Laboratories, CLM-1822-H) or U-^13^C-Glc (Cambridge Isotope Laboratories, CLM-1396), and the appropriate complementary substrate (unlabeled glutamine or glucose). Cells were plated 24 hours prior to labeling at 500,000 cells per well in 6-well plates. Cells were labeled overnight to achieve steady-state labeling. Metabolites were extracted and data was collected according to previously described procedures (ref:^22^). Data were processed as described in the unlabeled targeted metabolomics section.

#### Gas chromatography

Cells were cultured as described above. Metabolite extraction was performed as described^61^. Briefly, cells were lysed with dry-ice cold 80% methanol and metabolite extracts were then centrifuged at 20,000 g for 7 min at 4°C. Chloroform (stabilized with amylene) was added to each clarified supernatant. Phase separation was reached by centrifugation at 20,000 g for 15 min at 4°C. The aqueous phase was lyophilized using a SpeedVac concentrator, snap frozen in liquid nitrogen, and stored at −80°C for further processing. Samples were dissolved in 30 μl of 2% methoxyamine hydrochloride in pyridine (MOX) (Pierce, TS-45950) at 37°C for 1.5 hrs. Samples were derivatized by adding 45μl of N-methyl-N-(tert-butyldimethylsilyl)trifluoroacetamide (MBTSTFA) + 1% tert-butyldimethylchlorosilane (TBDMCS) (Pierce, TS-48927) at 60°C for 1 hr.

GC-MS analysis was performed as described^62^. Briefly, analysis was performed on an Agilent 6890 GC instrument that contained a 30m DB-35MS capillary column, which was interfaced to an Agilent 5975B MS. Electron impact (EI) ionization was set at 70 eV. Each analysis was operated in scanning mode, recording mass-to-charge-ratio spectra in the range of 100 – 605 m/z. For each sample 1μl was injected at 270°C, using helium as the carrier gas at a flow rate of 1 ml/min. To mobilize metabolites, the GC oven temperature was held at 100°C for 3 min and increased to 300°C at 3.5°C/min.

### Xenograft tumors and treatments

All animal studies were performed in accordance with the guidelines of Institutional Animal Care and Use Committee (IACUC) and approved protocols. NOD scid gamma (NSG) mice (Jackson Laboratory, 005557), 6-8 or 8-10 weeks old of both sexes, were maintained in the facilities of the Unit for Laboratory Animal Medicine (ULAM) under specific pathogen-free conditions. Mice were subcutaneously (s.c.) injected in both flanks with 0.5×10^6^ total cells (2.0×10^6^ for HCT 116) of iDox-shGOT1 #1 or shNT (n=8, iDox-shGOT1 BxPC-3 +/- dox, iDox-shNT BxPC-3 +dox tumors; n=6, iDox-shGOT1 PA-TU-8902 +/- dox, iDox-shNT PA-TU-8902 +/- dox, iDox-shNT BxPC-3-dox, iDox-shNT DLD-1 +/- dox tumors; n=5, iDox-shGOT1 HCT 116 +dox, iDox-shGOT1 DLD-1 +/-dox tumors; n=4 iDox-shGOT1 HCT 116-dox, iDox-shNT HCT 116 +/- dox tumors). Stable cells were trypsinized (Gibco, 25300-054) and suspended at 1:1 ratio of DMEM (Gibco, 11965-092) cell suspension:Matrigel (Corning, 354234) in 150-200 μL/injection. Dox chow (BioServ, F3949) was fed to the +dox groups on day 7 post tumor s.c. injection. Tumor size was assessed using a digital caliper twice/week after tumor cell implantation. Tumor volume (V, mm^3^) was calculated as V =1/2(length x width^2^) or V =π/6(length x width^2^) (ref:^63^). At endpoint, mice were sacrificed and final volume and mass of tumors were measured prior to tissue processing. Tissue was either snap-frozen in liquid nitrogen and stored at −80°C until processed for protein or metabolite analysis, or fixed in zinc formalin fixative (Z-Fix, Anatech LTD, #174) solution for >24 hours then replaced with 70% ethanol for future histological and/or histochemical staining

For radiotherapy studies, mice were randomized to receive no treatment (mock), dox alone (dox), radiation alone (rad), or combined treatment (dox+rad) (n=12 tumors per arm except n=10 dox HCT 116 tumors). Radiation (2 Gy/fraction) was administered over 6 daily fractions, beginning day 10 after implantation) using a Philips RT250 (Kimtron Medical) unit at a dose rate of approximately 2 Gy/minute. Dosimetry was performed using an ionization chamber directly traceable to a National Institute of Standards and Technology calibration. Animals were anesthetized with isoflurane and positioned such that the apex of each flank tumor was at the center of a 2.4 cm aperture in the secondary collimator, with the rest of the mouse shielded from radiation^60^.

### CCLE Dataset Analysis

The cancer cell line encyclopedia (CCLE) dataset with accession number GSE36133 (ref:^64^) was downloaded from the NCBI Gene Expression Omnibus^65^. The mRNA expression value of genes encoding GOT1-related enzymes, i.e. malic enzymes (ME1/2), malate dehydrogenases (MDH1/2), transaminases (GOT1/2, GPT1/2), and glutaminolysis enzymes (GLS, GLUD1) in PDA cell lines were compared to those of CRC.

### Seahorse Analysis

Extracellular acidification rates (ECAR) were performed using the XF-96 Extracellular Flux Analyzer (Agilent Technologies). Cells were treated with dox for four days then plated on Seahorse Microplates in DMEM media (+/-dox) at: PA-TU-8902 30,000 cells/well, PA-TU-8988T 60,000 cells/well, HCT 116 60,000 cells/well, DLD-1 60,000 cells/well. The next day, media was replaced with Seahorse XF Base DMEM (Agilent, 103335-100) containing 25 mM glucose and 2 mM glutamine adjusted to pH ~7.4, and the plate was allowed to incubate for 1 hour in a non-CO_2_, 37°C incubator. For the mitostress test, ECAR was measured under basal conditions and in response to mitochondrial inhibitors: oligomycin (0.5 μM), FCCP (0.25 μM), rotenone (0.5 μM), and antimycin A (0.5 μM).

### Genome-Scale Metabolic Network Modeling Using Dynamic Flux Analysis

PA-TU-8902 cells were plated in 6-well plates in triplicate for each timepoint (4, 2, or 0 days of dox treatment). Media and cells were collected separately for unlabeled targeted metabolomics. The metabolomics data were used as constraints in the human metabolic reconstruction^26^ to create a metabolic model using the dynamic flux analysis (DFA) approach^28,29^. DFA determines the optimal metabolic state that satisfies the biomass objective function and metabolomic constraints. DFA uses measured rate of change from time-course metabolomics data to constrain fluxes. Because subcellular compartment information is lost during metabolomics measurement, we assumed that the measured metabolites represent the sum total in the cytoplasm, nucleus, and mitochondrial compartments. We used both intracellular and extracellular metabolite measurements to construct the metabolic models. Single gene and reaction knockouts were conducted on the metabolic models to estimate their impact on cellular growth rate. These models were used to identify genes and metabolic reactions that were differentially active between +/- dox cells.

### Cell Viability Assay

PDA and CRC cells were plated at densities for log growth in the presence of dox (or mock treatment) for 4 days. On day 4, cells were trypsinized (Gibco, 25300-054) and replated in triplicate in 100 μL at 1,000 cells/well for-dox groups (PDA and CRC) and at 1,000 cells/well (CRC) or 3,000 cells/well (PDA) for +dox groups in white-walled 96 well plates (Corning/Costar, 3917). Cells were treated the following day with serial dilution of gemcitabine (Cayman Chemical, 9003096), 5-FU (Cayman Chemical, 14416), or oxaliplatin (Cayman Chemical, 13106). Cell viability was measured after 3 days using the CellTiter-Glo 2.0 Cell Viability Assay (Promega, G9243). Luminescence was measured for 500ms using a SpectraMax M3 Microplate Reader (Molecular Devices). IC50 values were calculated using GraphPad Prism 7 using three-parameter nonlinear regression analysis (except gemcitabine treated PA-TU-8902 used normalized response nonlinear regression analysis).

### Glutathione Enzymatic Assay

Cells were grown in +/- dox media for 3 days then plated in 96-well plates in +/-dox media. The following day, GSH/GSSG ratio was measured according to the manufacturer instructions (Promega, V6611).

### Statistical Analysis

Statistics were calculated using GraphPad Prism 7. One-way ANOVA was performed for experiments comparing multiple groups with one changing variable. ANOVA analyses were followed by Tukey’s post hoc tests to allow multiple group comparisons. A Student’s t-test (unpaired, two-tailed) was performed when comparing two groups to each other. Metabolomics data comparing 3 PDA and 3 CRC cell lines was analyzed by performing a Student’s t-test (unpaired, two-tailed) between all PDA metabolites and CRC metabolites. Time to tumor tripling analysis was performed using log-rank (Mantel–Cox) test. Outliers were removed with GraphPad using Grubbs’ test, alpha=0.05. Groups were considered significantly different when *P*< 0.05. All data are presented as mean ± s.d. (standard deviation).

