## Supplementary material for "Tissue of Origin Dictates GOT1 Dependence and Confers Synthetic Lethality to Radiotherapy": Nelson, et al_Extended Figures

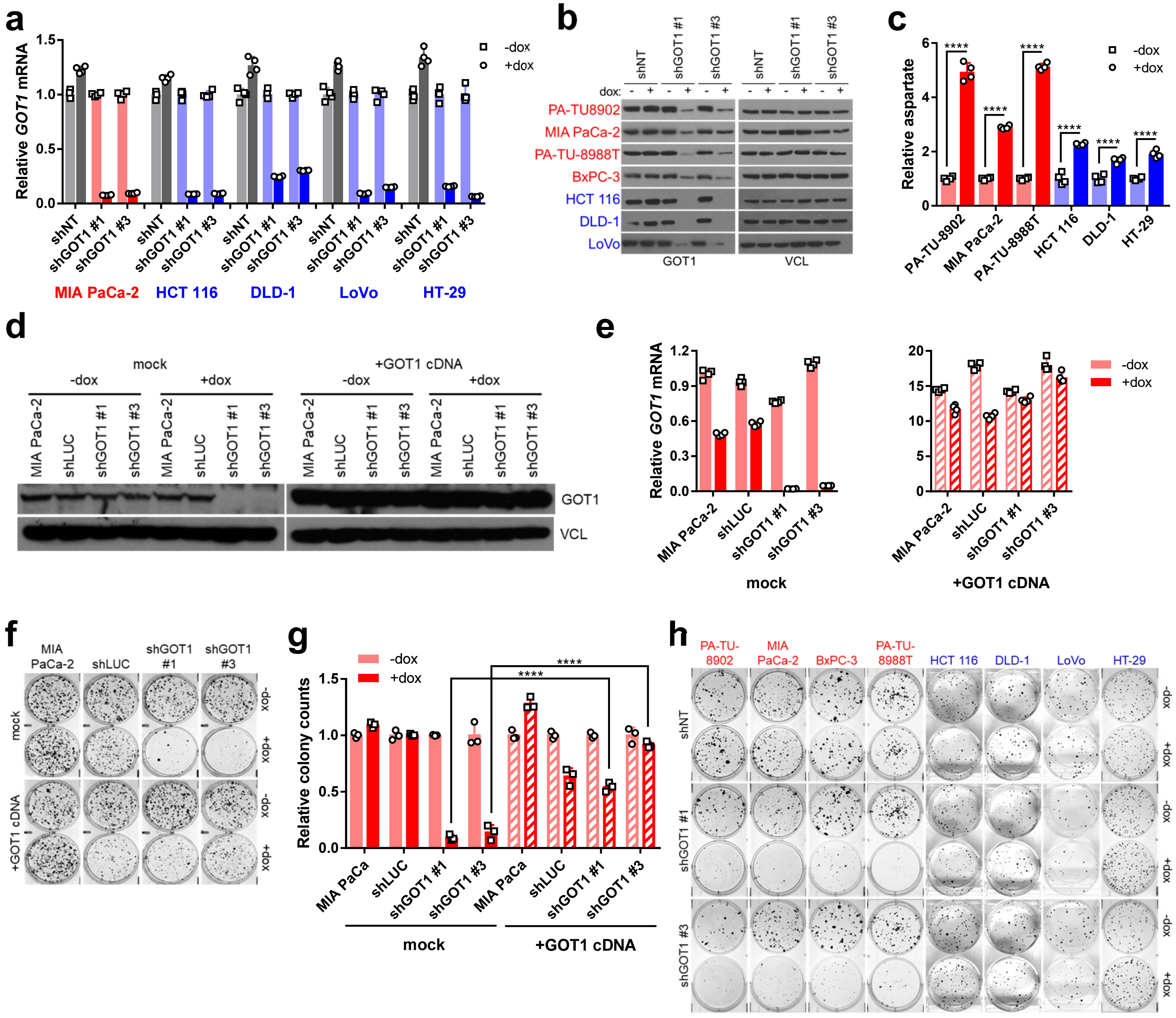

a
Relative GOT1 mRNA
-dox
+dox
shNT
shGOT1 #1
shGOT1 #3
MIA PaCa-2
HCT 116
DLD-1
LoVo
HT-29
b
dox:
PA-TU8902
MIA PaCa-2
PA-TU-8988T
BxPC-3
HCT 116
DLD-1
LoVo
GOT1
VCL
c
Relative aspartate
PA-TU-8902
MIA PaCa-2
PA-TU-8988T
HCT 116
DLD-1
HT-29
d
mock
+GOT1 cDNA
-dox
+dox
MIA PaCa-2
shLUC
shGOT1 #1
shGOT1 #3
GOT1
VCL
e
Relative GOT1 mRNA
mock
+GOT1 cDNA
f
g
Relative colony counts
MIA PaCa
h
PA-TU-8902
MIA PaCa-2
BxPC-3
PA-TU-8988T
HCT 116
DLD-1
LoVo
HT-29
shNT
shGOT1 #1
shGOT1 #3


E1

**Extended Figure 1: Description and validation of GOT1 knockdown system.** (**a**) *GOT1* mRNA expression, as determined by qPCR. Error bars represent s.d. from biological replicates (n=3). (**b**) Western blots for GOT1 and vinculin (VCL) loading control from iDox-shNT, iDox-shGOT1 #1, and iDox-shGOT1 #3 PDA and CRC cell lines +/- dox treatment. (**c**) Relative aspartate levels in iDox-shGOT1 PDA and CRC cell lines, as determined by LC/MS. Error bars represent s.d. from biological replicates (n=4). (**d**) Western blots for GOT1 and VCL from iDox-shGOT1 #1 and iDox-shGOT1 #3 MIA PaCa-2 PDA cells +/- dox treatment, +/- rescue with an ectopic shRNA-resistant GOT1 cDNA. (**e**) *GOT1* mRNA expression in MIA PaCa-2 PDA cells +/- dox treatment, +/- rescue with an ectopic shRNA-resistant GOT1 cDNA. Error bars represent s.d. from biological replicates (n=4). (**f**) Representative wells from colony forming assays and (**g**) associated quantitation in MIA PaCa-2 PDA cells +/- dox treatment, +/- GOT1 cDNA rescue. Error bars represent s.d. from biological replicates (n=3). (**h**) Representative wells from colony forming assays of cells expressing the iDox-shNT, iDox-shGOT1 #1, and iDox-shGOT1 #3 hairpins +/-dox. ****, $P < 0.0001$; Student's t-test (unpaired, two-tailed).

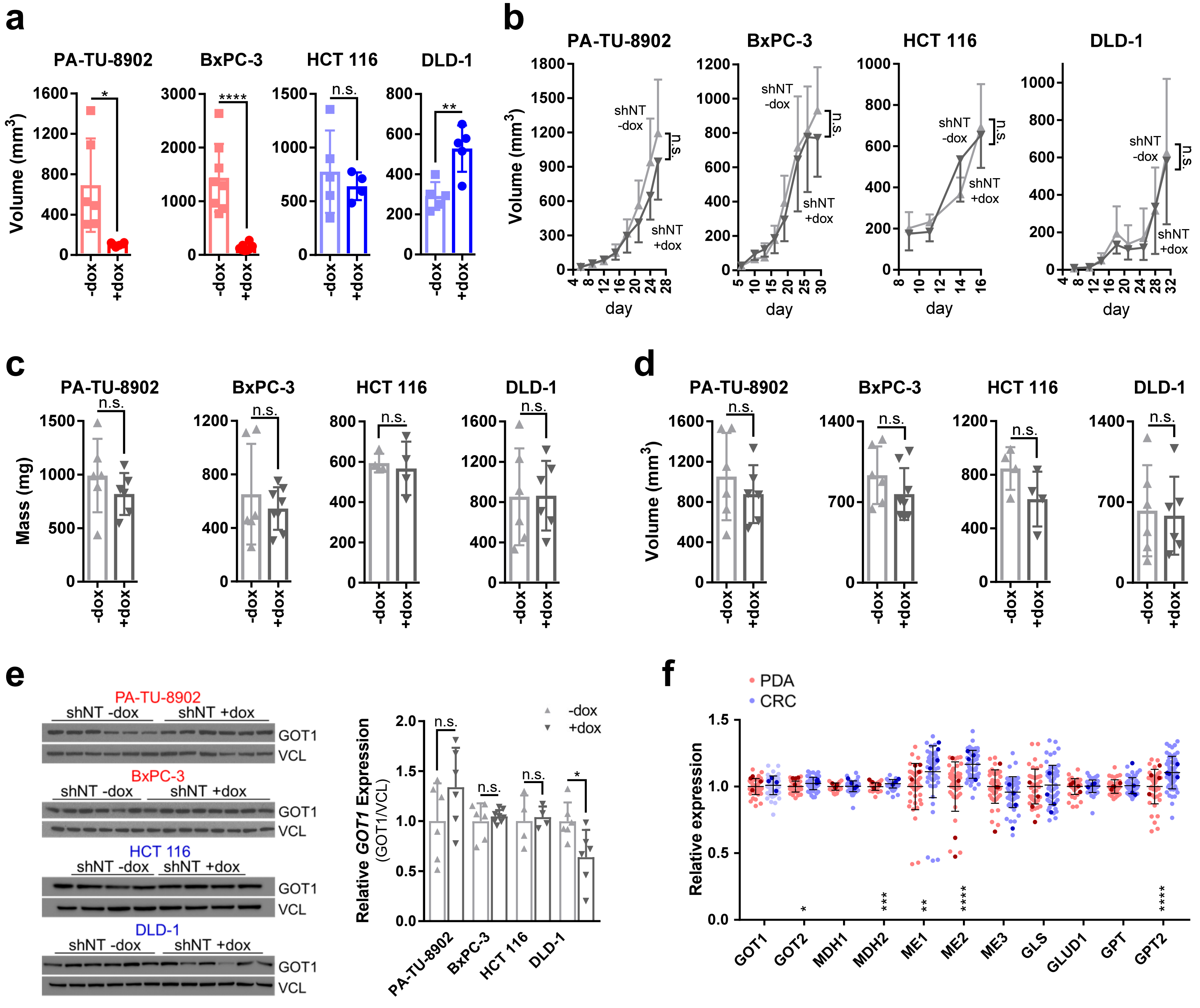

a
PA-TU-8902
BxPC-3
HCT 116
DLD-1
Volume (mm3)
-dox
+dox
n.s.
b
shNT -dox
shNT +dox
day
c
Mass (mg)
d
e
shNT -dox
shNT +dox
GOT1
VCL
Relative GOT1 Expression (GOT1/VCL)
f
PDA
CRC
Relative expression
GOT1
GOT2
MDH1
MDH2
ME1
ME2
ME3
GLS
GLUD1
GPT
GPT2


E2

**Extended Figure 2: *In vivo* tumor models of GOT1 knockdown and GOT1 pathway expression.** (**a**) Final tumor volume of PDA and CRC iDox-shGOT1 #1 lines in mice administered chow with or without dox (n=8, BxPC-3 +/-dox tumors; n=6, PA-TU-8902 +/- dox tumors; n=5, DLD-1 +/-dox, HCT 116 +dox tumors; n=4, HCT 116 -dox tumors). (**b**) Subcutaneous xenograft tumor growth, (**c**) final tumor mass, and (**d**) final tumor volume of PDA and CRC cells expressing iDox-shNT in mice administered chow with or without dox (n=8, BxPC-3 +dox tumors; n=6, BxPC-3 -dox, PA-TU-8902 +/- dox, DLD-1 +/-dox tumors; n=4 HCT 116 +/- dox tumors). (**e**) Western blots for GOT1 and VCL from iDox-shNT PDA and CRC tumors. (**f**) Relative mRNA expression level of the GOT1-pathway members, homologues, and adjacent components in PDA and CRC cells. Data obtained from the Cancer Cell Line Encyclopedia (CCLE)[1]. Data for the PDA and CRC cell lines used herein are highlighted in darker shades of red and blue, respectively. GLS, glutaminase; GLUD1, glutamate dehydrogenase 1; GOT1, glutamate oxaloacetate transaminase 1; GOT2, glutamate oxaloacetate transaminase 2; GPT, glutamate pyruvate transaminase; GPT2, glutamate pyruvate transaminase 2; MDH1, malate dehydrogenase 1; MDH2, malate dehydrogenase 2; ME1, malic enzyme 1; ME2, malic enzyme 2; ME3, malic enzyme 3. n.s., not significant; *, $P < 0.05$; **, $P < 0.01$; ***, $P < 0.001$; ****, $P < 0.0001$; Student's t-test (unpaired, two-tailed).

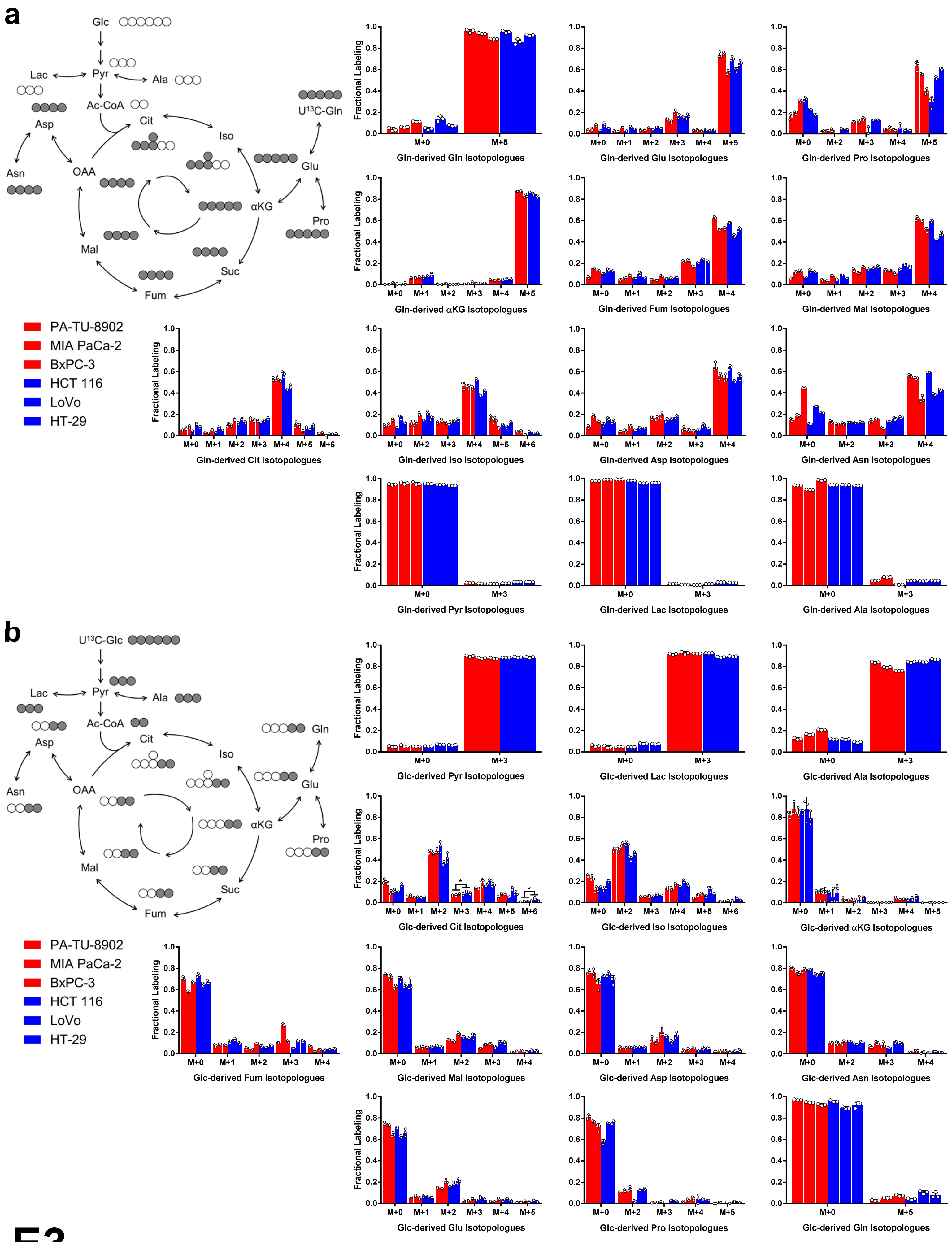
a
Glc
Lac
Pyr
Ala
Ac-CoA
Asp
Cit
U13C-Gln
Iso
Asn
OAA
Glu
αKG
Pro
Mal
Suc
Fum
PA-TU-8902
MIA PaCa-2
BxPC-3
HCT 116
LoVo
HT-29
Fractional Labeling
Gln-derived Gln Isotopologues
Gln-derived Glu Isotopologues
Gln-derived Pro Isotopologues
Gln-derived αKG Isotopologues
Gln-derived Fum Isotopologues
Gln-derived Mal Isotopologues
Gln-derived Cit Isotopologues
Gln-derived Iso Isotopologues
Gln-derived Asp Isotopologues
Gln-derived Asn Isotopologues
Gln-derived Pyr Isotopologues
Gln-derived Lac Isotopologues
Gln-derived Ala Isotopologues
b
U13C-Glc
Gln
Glc-derived Pyr Isotopologues
Glc-derived Lac Isotopologues
Glc-derived Ala Isotopologues
Glc-derived Cit Isotopologues
Glc-derived Iso Isotopologues
Glc-derived αKG Isotopologues
Glc-derived Fum Isotopologues
Glc-derived Mal Isotopologues
Glc-derived Asp Isotopologues
Glc-derived Asn Isotopologues
Glc-derived Glu Isotopologues
Glc-derived Pro Isotopologues
Glc-derived Gln Isotopologues

E3

**Extended Figure 3: Steady state fractional labeling patterns of TCA cycle and branching metabolites from glucose and glutamine carbon tracing in PDA and CRC cell lines.** Fractional labeling patterns reflect the percentage of a given metabolite pool labeled by an input metabolic substrate (in this case, glucose or glutamine carbon). (**a**) Isotopologue distributions presented as fractional enrichment from overnight labeling with $^{13}C$-glutamine (Gln) and (**b**) $^{13}C$-glucose (Glc) in PDA (red) and CRC (blue) cell lines, as determined by LC/MS, except for pyruvate, lactate, alanine, asparagine, and proline, which were generated by gas chromatography (GC)/MS. In the schemes at top left, filled circles represent $^{13}C$-labeled carbon, and open circles represent unlabeled carbon. The labeling pattern for one turn of the TCA cycle is presented. Citrate data previously published in a methods paper[2]. Error bars represent s.d. from biological replicates (n=3). Ac-CoA, acetyl-CoA; αKG, alpha-ketoglutarate; Ala, alanine; Asn, asparagine; Asp, aspartate; Cit, citrate; Fum, fumarate; Glu, glutamate; Iso, isocitrate; Lac, lactate; Mal, malate; OAA, oxaloacetate; Pro, proline; Pyr, pyruvate; $U^{13}C$, uniformly labeled carbon.

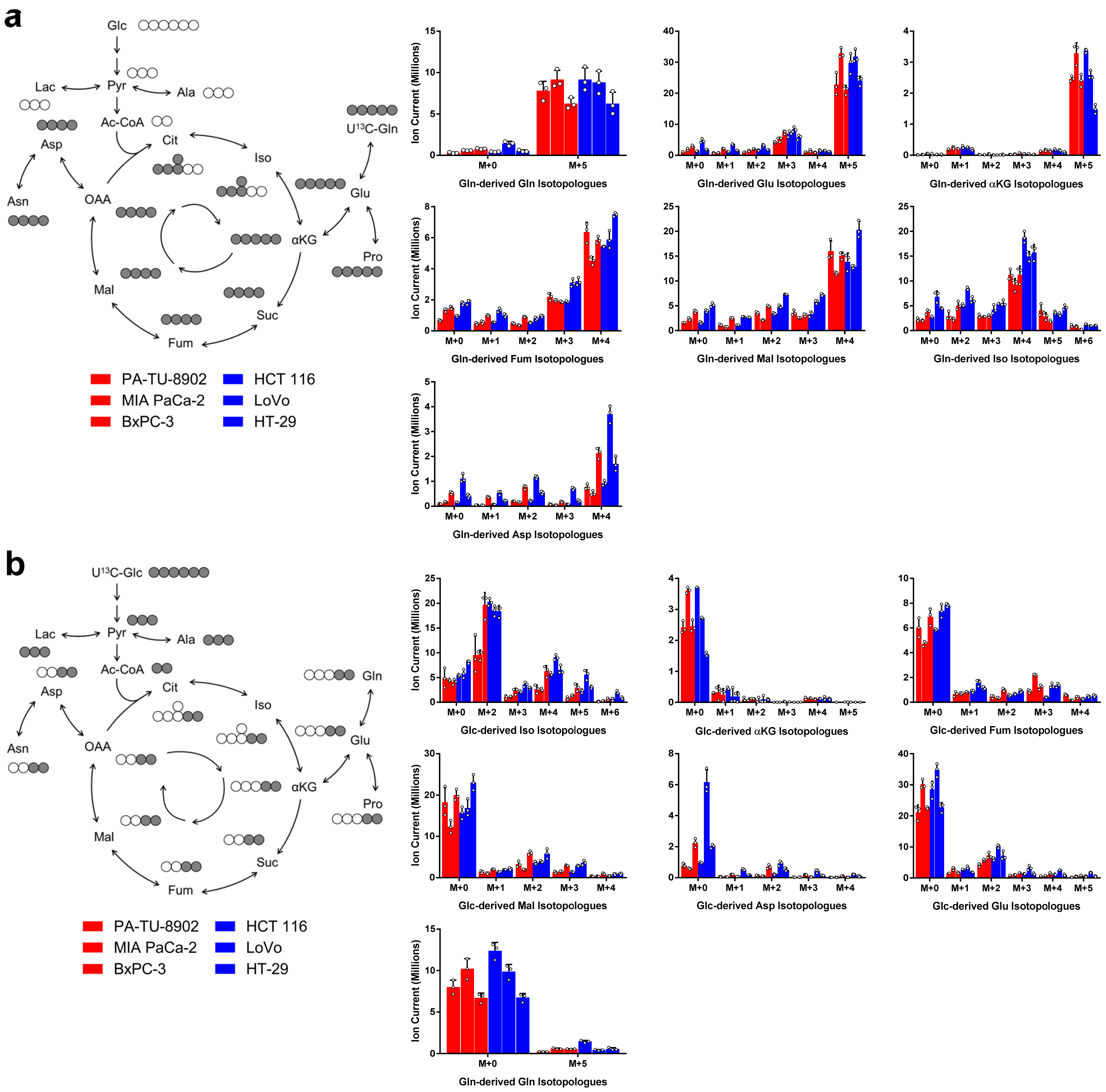
a
Glc
Lac
Pyr
Ala
Ac-CoA
Asp
Cit
U13C-Gln
Iso
Asn
OAA
Glu
αKG
Pro
Mal
Suc
Fum
PA-TU-8902
MIA PaCa-2
BxPC-3
HCT 116
LoVo
HT-29
Ion Current (Millions)
Gln-derived Gln Isotopologues
Gln-derived Glu Isotopologues
Gln-derived αKG Isotopologues
Gln-derived Fum Isotopologues
Gln-derived Mal Isotopologues
Gln-derived Iso Isotopologues
Gln-derived Asp Isotopologues
b
U13C-Glc
Gln
Glc-derived Iso Isotopologues
Glc-derived αKG Isotopologues
Glc-derived Fum Isotopologues
Glc-derived Mal Isotopologues
Glc-derived Asp Isotopologues
Glc-derived Glu Isotopologues
Gln-derived Gln Isotopologues

# E4

**Extended Figure 4: Steady state pools of TCA cycle and branching metabolites from glucose and glutamine carbon tracing in PDA and CRC cell lines.** Pool sizes reflect the relative abundance of a given metabolite. Herein, these are presented as the total of labeled and unlabeled metabolite. (**a**) Isotopologue distributions presented as total metabolite pools from overnight labeling with $^{13}C$-glutamine (Gln) and (**b**) $^{13}C$-glucose (Glc) in PDA (red) and CRC (blue) cell lines, as determined by LC/MS. In the schemes at top left, filled circles represent $^{13}C$-labeled carbon, and open circles represent unlabeled carbon. Labeling pattern for one turn of the TCA cycle is presented. Citrate data previously published in a methods paper[2]. Error bars represent s.d. from biological replicates (n=3). Ac-CoA, acetyl-CoA; αKG, alpha-ketoglutarate; Ala, alanine; Asn, asparagine; Asp, aspartate; Cit, citrate; Fum, fumarate; Glu, glutamate; Iso, isocitrate; Lac, lactate; Mal, malate; OAA, oxaloacetate; Pro, proline; Pyr, pyruvate; $U^{13}C$, uniformly labeled carbon.

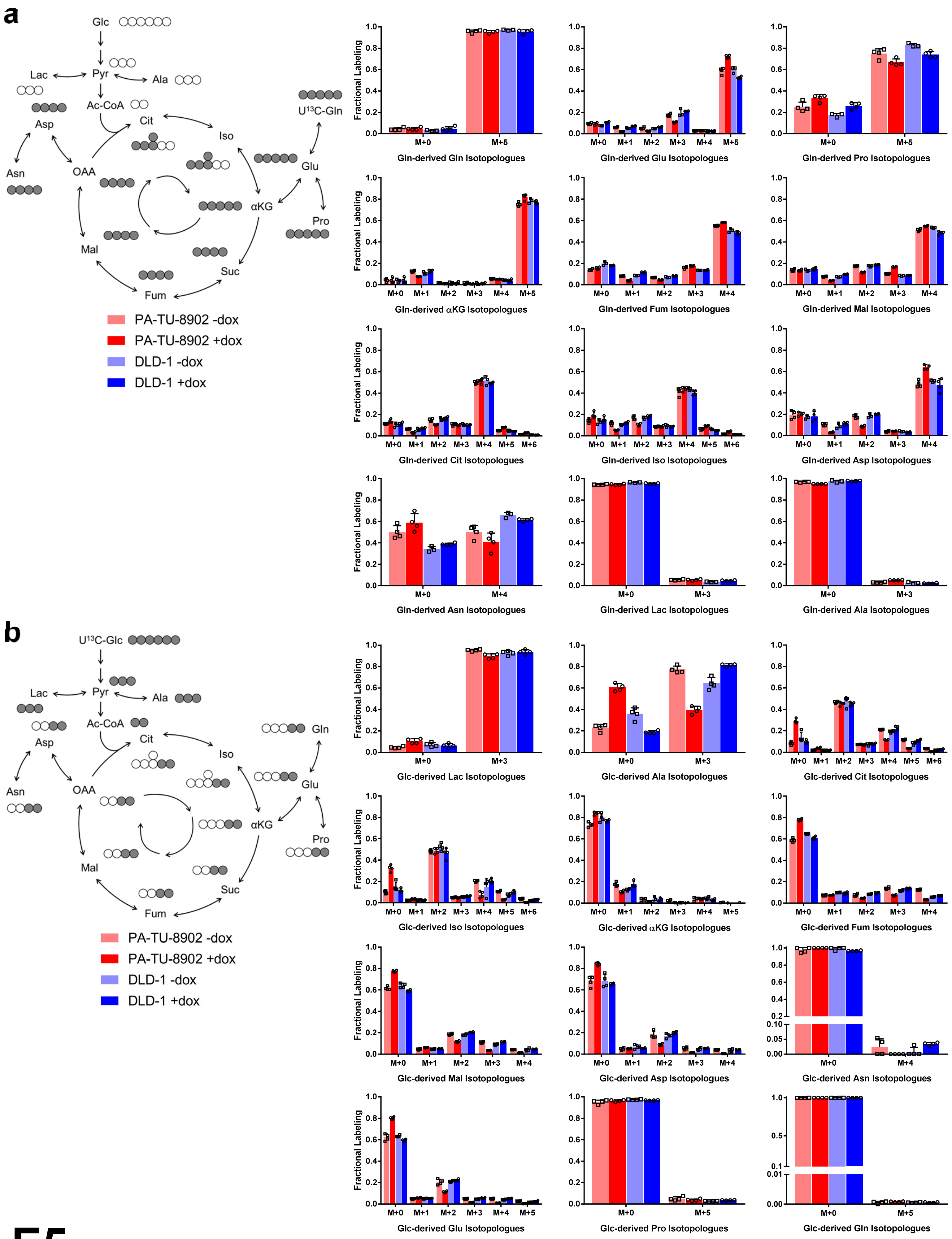

a
Glc
Lac
Pyr
Ala
Ac-CoA
Asp
Cit
Iso
U13C-Gln
Asn
OAA
Glu
αKG
Pro
Mal
Suc
Fum
PA-TU-8902 -dox
PA-TU-8902 +dox
DLD-1 -dox
DLD-1 +dox
Fractional Labeling
Gln-derived Gln Isotopologues
Gln-derived Glu Isotopologues
Gln-derived Pro Isotopologues
Gln-derived αKG Isotopologues
Gln-derived Fum Isotopologues
Gln-derived Mal Isotopologues
Gln-derived Cit Isotopologues
Gln-derived Iso Isotopologues
Gln-derived Asp Isotopologues
Gln-derived Asn Isotopologues
Gln-derived Lac Isotopologues
Gln-derived Ala Isotopologues
b
U13C-Glc
Gln
Glc-derived Lac Isotopologues
Glc-derived Ala Isotopologues
Glc-derived Cit Isotopologues
Glc-derived Iso Isotopologues
Glc-derived αKG Isotopologues
Glc-derived Fum Isotopologues
Glc-derived Mal Isotopologues
Glc-derived Asp Isotopologues
Glc-derived Asn Isotopologues
Glc-derived Glu Isotopologues
Glc-derived Pro Isotopologues
Glc-derived Gln Isotopologues


E5

**Extended Figure 5: Steady state fractional labeling patterns of TCA cycle and branching metabolites from glucose and glutamine carbon tracing in PDA and CRC after GOT1 knockdown.** (**a**) Isotopologue distributions presented as fractional enrichment from overnight labeling with $^{13}C$-glutamine (Gln) and (**b**) $^{13}C$-glucose (Glc) in iDox-shGOT1 #1 PA-TU-8902 PDA (red) and DLD-1 CRC (blue) cell lines. GOT1 was knocked down for 5 days via dox treatment; metabolites patterns determined by LC/MS. In the schemes at top left, filled circles represent $^{13}C$-labeled carbon, and open circles represent unlabeled carbon. Labeling pattern for one turn of the TCA cycle is presented. Error bars represent s.d. from biological replicates (n=4, except n=3 for DLD-1 -dox Gln labeling). Ac-CoA, acetyl-CoA; αKG, alpha-ketoglutarate; Ala, alanine; Asn, asparagine; Asp, aspartate; Cit, citrate; Fum, fumarate; Glu, glutamate; Iso, isocitrate; Lac, lactate; Mal, malate; OAA, oxaloacetate; Pro, proline; Pyr, pyruvate; $U^{13}C$, uniformly labeled carbon.

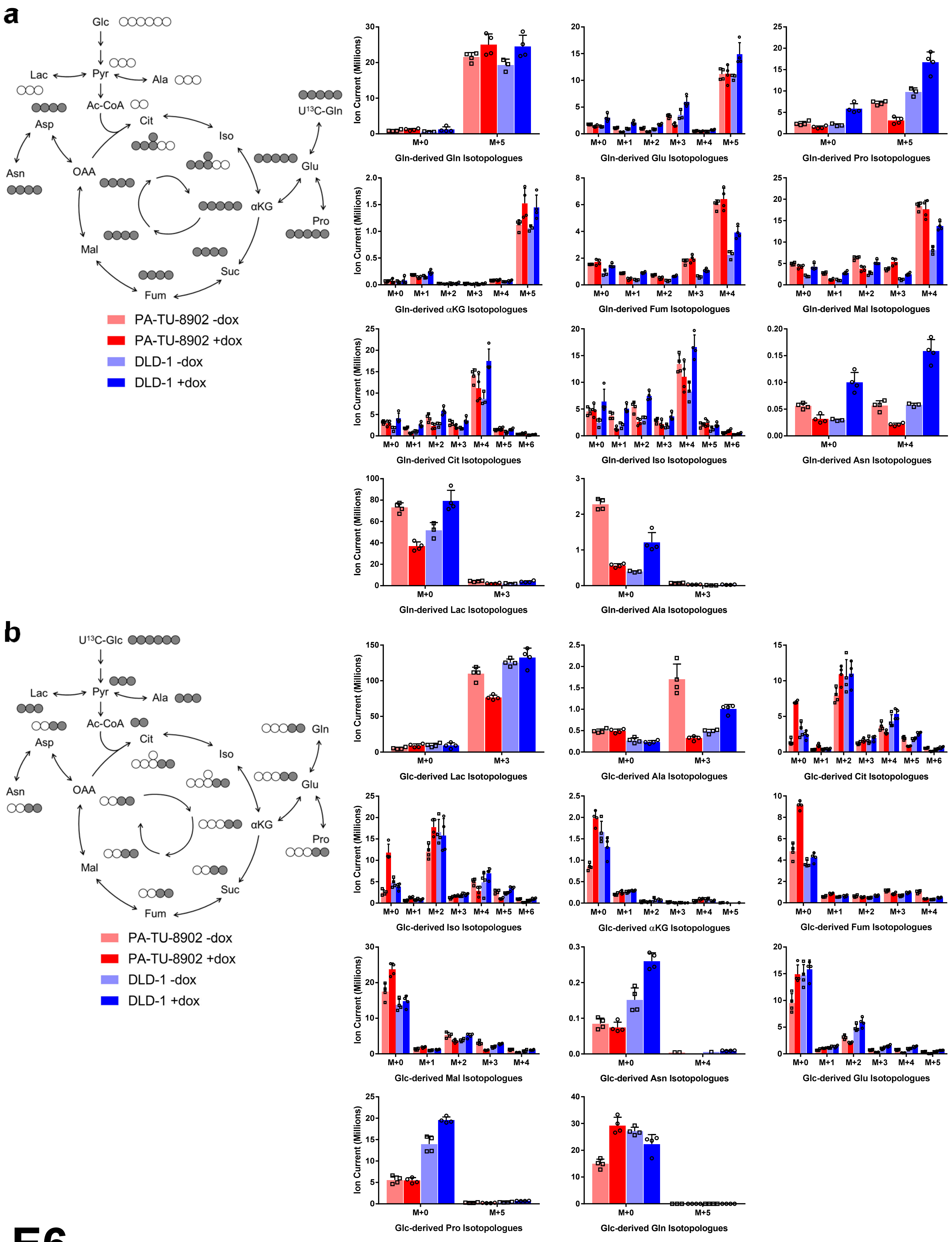
a
Glc
Lac
Pyr
Ala
Ac-CoA
Asp
Cit
Iso
U¹³C-Gln
Asn
OAA
Glu
αKG
Pro
Mal
Suc
Fum
PA-TU-8902 -dox
PA-TU-8902 +dox
DLD-1 -dox
DLD-1 +dox
Ion Current (Millions)
Gln-derived Gln Isotopologues
Gln-derived Glu Isotopologues
Gln-derived Pro Isotopologues
Gln-derived αKG Isotopologues
Gln-derived Fum Isotopologues
Gln-derived Mal Isotopologues
Gln-derived Cit Isotopologues
Gln-derived Iso Isotopologues
Gln-derived Asn Isotopologues
Gln-derived Lac Isotopologues
Gln-derived Ala Isotopologues
b
U¹³C-Glc
Lac
Pyr
Ala
Ac-CoA
Asp
Cit
Gln
Iso
Asn
OAA
Glu
αKG
Pro
Mal
Suc
Fum
PA-TU-8902 -dox
PA-TU-8902 +dox
DLD-1 -dox
DLD-1 +dox
Glc-derived Lac Isotopologues
Glc-derived Ala Isotopologues
Glc-derived Cit Isotopologues
Glc-derived Iso Isotopologues
Glc-derived αKG Isotopologues
Glc-derived Fum Isotopologues
Glc-derived Mal Isotopologues
Glc-derived Asn Isotopologues
Glc-derived Glu Isotopologues
Glc-derived Pro Isotopologues
Glc-derived Gln Isotopologues

E6

**Extended Figure 6: Steady state pools of TCA cycle and branching metabolites from glucose and glutamine carbon tracing in PDA and CRC after GOT1 knockdown.** (**a**) Isotopologue distributions presented as total metabolite pools from overnight labeling with $^{13}C$-glutamine (Gln) and (**b**) $^{13}C$-glucose (Glc) in iDox-shGOT1 #1 PA-TU-8902 PDA (red) and DLD-1 CRC (blue) cell lines. GOT1 was knocked down for 5 days via dox treatment; metabolites patterns determined by LC/MS. In the schemes at top left, filled circles represent $^{13}C$-labeled carbon, and open circles represent unlabeled carbon. Labeling pattern for one turn of the TCA cycle is presented. Error bars represent s.d. from biological replicates (n=4, except n=3 for DLD-1 -dox Gln labeling). Ac-CoA, acetyl-CoA; αKG, alpha-ketoglutarate; Ala, alanine; Asn, asparagine; Asp, aspartate; Cit, citrate; Fum, fumarate; Glu, glutamate; Iso, isocitrate; Lac, lactate; Mal, malate; OAA, oxaloacetate; Pro, proline; Pyr, pyruvate; $U^{13}C$, uniformly labeled carbon.

**a**

**Pathways Significantly Enriched upon GOT1 Knockdown in Vitro**

| Pathway Name | Hits | p | FDR |
|---|---|---|---|
| Pyrimidine metabolism | 9/60 | 1.31E-06 | 1.05E-04 |
| Purine metabolism | 10/92 | 6.56E-06 | 2.62E-04 |
| Cysteine and methionine metabolism | 6/56 | 6.21E-04 | 0.0166 |
| Sulfur metabolism | 3/18 | 0.0046 | 0.0927 |
| Nitrogen metabolism | 4/39 | 0.0064 | 0.1018 |
| Nicotinate and nicotinamide metabolism | 4/44 | 0.0098 | 0.1304 |
| Glycine, serine and threonine metabolism | 4/48 | 0.0132 | 0.1402 |
| Pantothenate and CoA biosynthesis | 3/27 | 0.0147 | 0.1402 |
| Arginine and proline metabolism | 5/77 | 0.0158 | 0.1402 |
| Glutathione metabolism | 3/38 | 0.0365 | 0.2602 |
| Cyanoamino acid metabolism | 2/16 | 0.0376 | 0.2602 |
| Glycerophospholipid metabolism | 3/39 | 0.0390 | 0.2602 |

**b**

**Pathways Significantly Enriched upon GOT1 Knockdown in Vivo**

| Pathway Name | Hits | p | FDR |
|---|---|---|---|
| Nicotinate and nicotinamide metabolism | 5/44 | 2.26E-04 | 0.0181 |
| Purine metabolism | 6/92 | 0.0011 | 0.0430 |
| Cysteine and methionine metabolism | 4/56 | 0.0058 | 0.1553 |
| Nitrogen metabolism | 3/39 | 0.0141 | 0.2583 |
| Aminoacyl-tRNA biosynthesis | 4/75 | 0.0161 | 0.2583 |
| Sulfur metabolism | 2/18 | 0.0229 | 0.3060 |
| Thiamine metabolism | 2/24 | 0.0394 | 0.4346 |
| Pyrimidine metabolism | 3/60 | 0.0438 | 0.4346 |
| Phenylalanine, tyrosine and tryptophan biosynthesis | 2/27 | 0.0489 | 0.4346 |

**c**

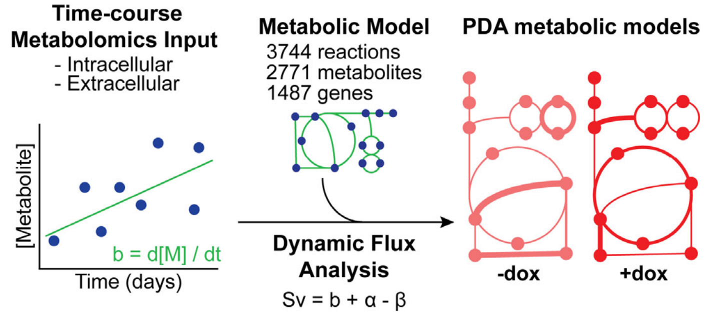


**d**

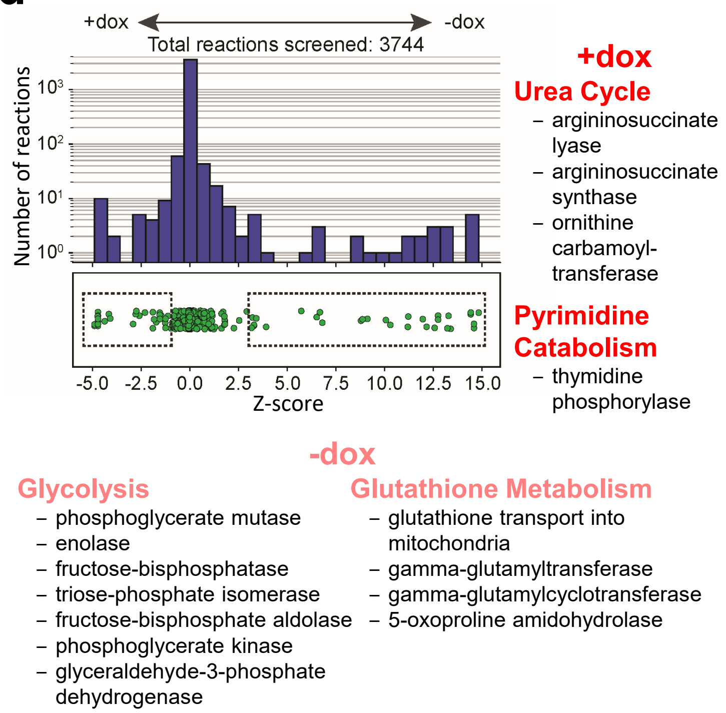


**e**

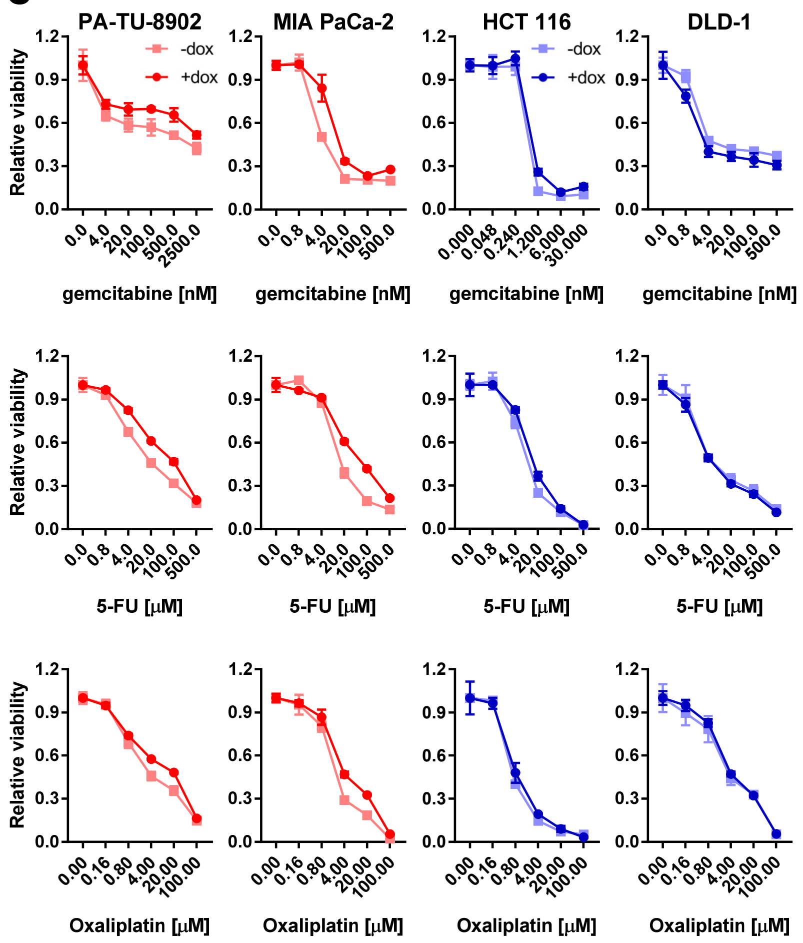

**Extended Figure 7: Metabolic pathways and drug responses associated with GOT1 inhibition.** (**a**) Pathway Analysis of metabolites from **Fig. 4a** with greater than 2-fold change, as determined using the Metaboanalyst web tool (https://www.metaboanalyst.ca/). Hits reflects number of metabolites that were significant over total of metabolites considered for a given pathway. FDR, false discovery rate. (**b**) Pathway Analysis via Metaboanalyst of metabolites determined by LC/MS with greater than 2-fold change from PDA PA-TU-8902 and CRC HCT 116 tumors from **Fig 1d,h**. Pathways highlighted in red were observed *in vitro* and *in vivo*. (**c**) Schematic overview of Dynamic Flux Analysis (DFA) approach for constructing genome-scale metabolic models of timecourse intracellular and extracellular metabolites from iDox-shGOT1 #1 PA-TU-8902 cells (+/- dox) as determined by LC/MS. In the schemes, circles represent metabolites connected in a metabolic pathway. DFA uses 3744 reactions, 2771 metabolites and 1487 genes. The network represented as the stoichiometric matrix (S) and is used to solve the reaction flux vector (v), determined by change of metabolite concentration over time (d[M] / dt or b). See Methods for further description. (**d**) Systematic reaction knockout analysis using DFA identifies differentially active metabolic reactions in PDA +/- GOT1 knockdown. “Knocked-out” reactions with negative Z-scores resulted in a computationally predicted decrease in PDA “growth” in cells under +dox conditions, while reactions with positive Z-scores decreased PDA “growth” in cells under -dox conditions. The y-axis shows the total number of reactions in each Z-score bin. Selected KEGG pathways from +/- dox DFA with related enzymes that reached statistical significance are shown to the right and bottom. Reactions shown have $P < 0.05$ (two-sample t-test). (**e**) Relative viability of iDox-shGOT1 #1 PDA and CRC cells treated with a dose response of gemcitabine (top), 5-fluorouracil (middle), and oxaliplatin (bottom). GOT1 was knocked down for 5 days before drug treatment, and +dox were maintained under dox for the duration of the experiment. Relative viability was calculated by cell titer glo at day 3. Error bars represent s.d. from biological replicates (n=3).

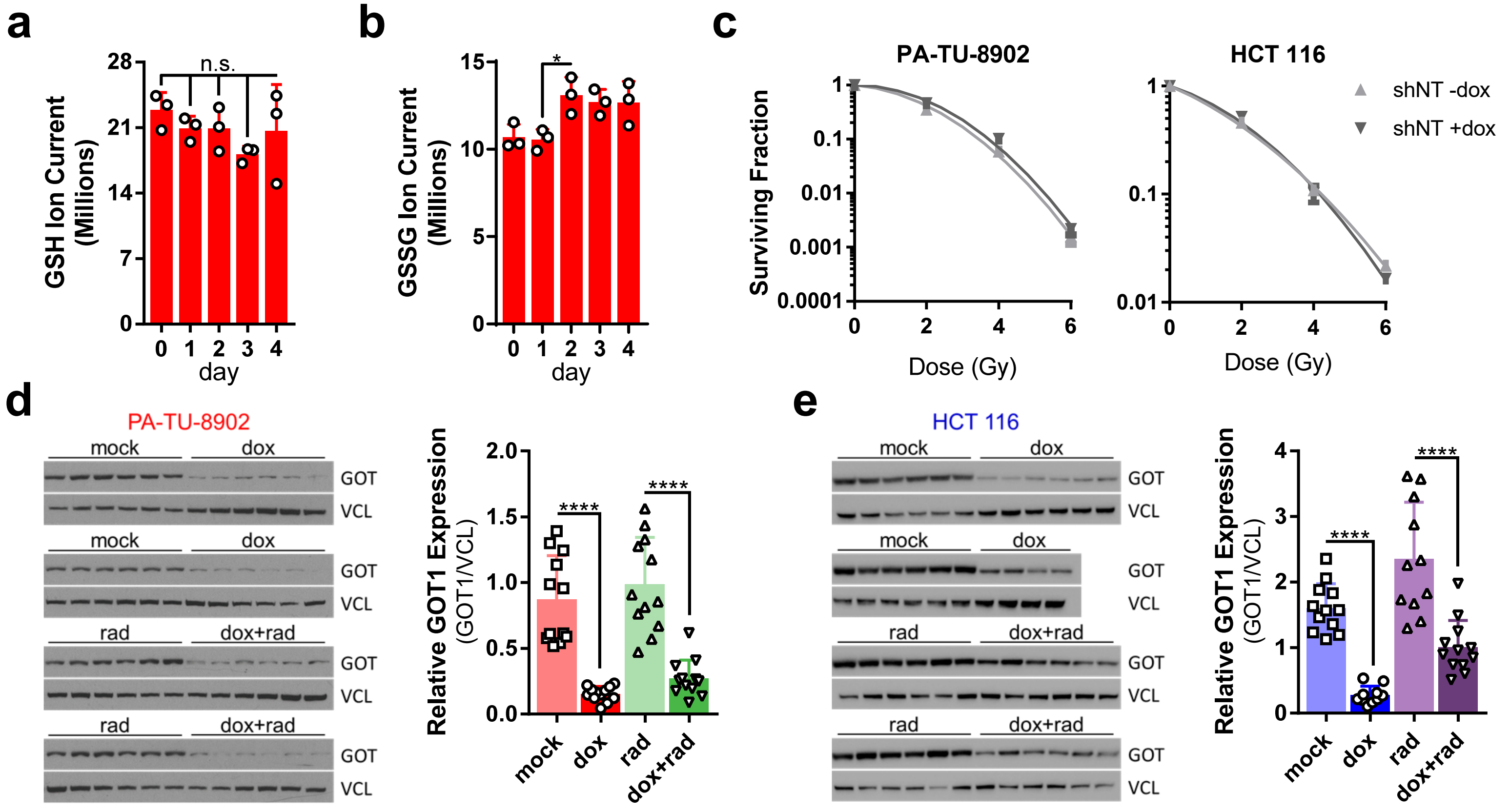
a
GSH Ion Current (Millions)
n.s.
day
b
GSSG Ion Current (Millions)
*
day
c
PA-TU-8902
HCT 116
Surviving Fraction
Dose (Gy)
shNT -dox
shNT +dox
d
PA-TU-8902
mock
dox
rad
dox+rad
GOT
VCL
Relative GOT1 Expression (GOT1/VCL)
****
e
HCT 116

**Extended Figure 8: Ancillary data associated with Fig 5.** Timecourse of total (**a**) reduced glutathione (GSH) and (**b**) oxidized glutathione (GSSG) pools in iDox-shGOT1 #1 PA-TU-8902 cells, as determined by LC/MS. (**c**) Surviving fraction from clonogenic assay of radiation-treated iDox-shNT PA-TU-8902 (left) and HCT116 (right), +/- dox. Error bars represent s.d. from biological replicates in **a-c** (n=3). Gy, Gray. Western blot (left) and quantification (right) for GOT1 expression in iDox-shGOT1 #1 (**d**) PA-TU-8902 and (**e**) HCT116 xenograft tumors treated with dox and/or radiation (rad) presented in **Fig. 5i-l**. Error bars represent s.d from n=12 tumors per arm except n=10 dox HCT 116 tumors. n.s., not significant; *, $P < 0.05$; ****, $P < 0.0001$; one-way ANOVA (**a,b**); Student's t-test (unpaired, two-tailed) (**d,e**).
